# HydroMEA: A 3D Hydrogel Based Microfluidic Device to Study Electrophysiology for Myelinated Nerve-on-Chip

**DOI:** 10.1101/2025.07.24.666400

**Authors:** Blandine F. Clément, Cédric Pfister, Timothy Kurer, Céline Labouesse, Dhanajay V. Deshmukh, Julian Hengsteler, Julia Lehmann, Lorenza G. Paganella, Tobias Ruff, Vilius Dranseika, Sean Weaver, Lukas Sommer, Mark W. Tibbitt, János Vörös, Christina M. Tringides

## Abstract

Engineered *in vitro* platforms are powerful systems to study information flow in the nervous system. While existing polydimethylsiloxane (PDMS)-based microfluidic platforms offer precise architectures, the cultured neurons grow on two-dimensional (2D) planar multielectrode arrays (MEA). To mimic the native microenvironment, where neurons grow in three-dimensional (3D) extracellular matrices (ECM), 3D hydrogels can be designed to encapsulate cells and enable physiologically-mimicked behaviors. Here, we describe ‘hydroMEA’, a 3D platform fabricated by placing PDMS microstructures on a high-density MEA and filled with a desired hydrogel, to offer controlled topologies, physiologically-relevant microenvironments, and real-time electrophysiological measurements. First, we developed a gelatin methacryloyl (GelMA) hydrogel with incorporated ECM components and tuned the mechanical properties to match those of nerve tissue. The hydrogel was able to support: 1) the growth of iPSC-derived sensory neurons (hSNs) for >100 days; 2) co-cultures of hSN with human embryonic stem cell-derived Schwann cells (hSCs), to enable reliable 3D myelination. Next, hydroMEA were prepared for topologically-defined 3D growth and myelination in designated compartments. Finally, electrophysiological evaluation of hSN-hSCs co-cultures revealed increased conduction speeds indicating functional myelin. This platform is a promising tool to study cell-cell interactions and to functionally evaluate the effect of pharmacological compounds in a more translational manner.

## 1. INTRODUCTION

In order to gain insights into neural development and neurological pathologies, it is critical to understand mechanisms of the nervous system. Electrophysiological observation of the nervous system advances our knowledge through the study of electrical information flow^1,2^. While advances in implantable neurotechnologies have enabled the monitoring of *in vivo* electrophysiology, these approaches face inherent limitations. These limitations compromise the ability to study detailed neural mechanisms, particularly those involving complex interactions between different cell types. For example, the formation and function of myelin, a fatty lipid sheath formed by Schwann cells wrapping around neuronal axons in the peripheral nervous system, is difficult to electrically monitor *in vivo* and is thus more readily evaluated with optical techniques^3^. Myelin not only facilitates the electrical axonal conduction, but is also a key modulator of disease development in the peripheral nervous system^4,5^.

Given the constraints of *in vivo* electrophysiology, there is a need for physiologically-relevant *in vitro* platforms that can recapitulate the complex structure and function of the nervous system, while enabling detailed mechanistic studies^6,7^. *In vitro* models and microphysiological systems (MPS) have emerged as powerful tools, as they offer controlled environments to study specific cellular interactions and molecular mechanisms^8–12^ while reducing the need for animal testing^13^. Beyond the investigation of fundamental mechanisms that govern neuronal function, physiologically-relevant *in vitro* platforms would have significant implications for the pharmaceutical industry. Progress in neurotherapeutic development continues to be hindered by the limited translatability of preclinical models, which rely on overly-simplified, non-physiological two-dimensional (2D) *in vitro* systems. This challenge highlights the urgent needs for MPS that can more accurately mimic cellular responses with human-derived cells, in platforms that can accurately recapitulate the complex functional and structural aspects of neural ECM, while maintaining experimental accessibility and reproducibility.

One strength of *in vitro* platforms is the reliable integration of the desired cell type(s) and co-cultures, and the ability to use human-derived cells. In the peripheral nervous system, a network of sensory and motor neurons connect to other organs and relay information to and from the central nervous system through a complex highway of nerves. To facilitate the flow of information, various nerves are myelinated by Schwann cells to increase conduction velocities and more quickly deliver neural signals. The process and mechanism(s) of myelination are not well understood, and require a co-culture of dorsal root ganglion (DRG) neurons and Schwann cells. However, most platforms combined rodent-derived DRG neurons with human-derived Schwann cells (hSCs)^14,15^, which have a reduced biological relevance. More recently, the emergence of human induced pluripotent stem cell (iPSC)-derived neuronal cells, including hiPSC-derived DRG or sensory neurons (hSNs), have further enhanced the translational potential of *in vitro* models. The successful co-culture of iPSC neurons with Schwann cells has supported the identification of improved culture conditions with some myelination reported^16,17^, and has served as a platform to investigate initial interactions between neurons and Schwann cells^18^. However, the absence of a 3D ECM-mimicked microenvironment is thought to contribute to the low-efficiency of myelination in 2D platforms^19^.

Neurons, *in vivo,* rely heavily on their mechanical microenvironment for cellular morphology, differentiation, and functionality^20,21^. Additionally, the mechanical properties of the cellular microenvironment, particularly the substrate stiffness and viscoelasticity, significantly modulate neuronal cell behavior including neurite branching patterns^22–24^, cell type differentiation, and network formation^25,26^. To match the architecture of ECM, various groups use electrospinning of polymers and incorporate SCs^27^ or human iPSC-derived cell sources^28,29^ into the scaffolds. However, the mechanical stiffness of these formulations is much higher than the properties of neural tissues. In contrast, hydrogels have emerged as materials that provide a 3D microenvironment with the same mechanical properties as the native neural mechanical microenvironment, and support cellular growth^30^. When hydrogels were integrated into *in vitro* platforms with a co-culture of neurons and myelinating glial cells, there was increased myelin formation for neurons grown in hydrogels as compared to neurons cultured on 2D substrates^18,26^. An added strength of hydrogels is the tunability of their chemical formulations. While Matrigel^TM^ has demonstrated success in neuronal cultures, it has significant limitations which include batch-to-batch variability, limited tunability of mechanical properties, and temperature-sensitivity^31,32^. There is increasing interest in naturally-derived and synthetic formulations, which have gained increasing attention for neural applications due to their mechanical tunability, versatility, and biocompatibility^33,34^. While these 3D models achieve greater biological relevance by combining human cells and 3D architectures, they lack precise spatial control and offer limited electrophysiological assessment capabilities.

In addition to the importance of mimicking the *in vivo* microenvironment, there is a need to combine qualitative imaging techniques with quantitative electrophysiology tools to comprehensively evaluate cell culture growth and behavior. Microelectrode array (MEA) techniques provide a powerful method to investigate neuronal network electrophysiology, in real-time, through underlying electrodes^35–37^. However, interfacing 3D neural tissues with flat MEA technologies remains a challenge. The 3D hydrogel or tissue that sits between cells and the underlying MEA hinder sufficient electrical coupling. This has driven the development of creative alternative strategies for electrophysiological measurements within 3D platforms including protruding electrode arrays^38^, improved tissue-MEA interactions by insulative compression^39^, or trapped microelectrodes for 3D neurites^40^. Despite these strategies, electrophysiological assessment of 3D neural cultures remains and ongoing challenge, specifically to address limitations in spatial accessibility, signal-to-noise ratio, and recording throughput.

In parallel, there have been increased efforts in achieving controlled topologies and guided neural growth to enhance signal read-out and build nerve-on-chip models of the peripheral nervous system. Microfluidic-based platforms, with features on the 10s-100s µm scale, have been developed to grow 2D neural cultures for drug screening^41,42,46^ or for rapid assessment of neural conduction speed in explanted nerve fibers^43,44^. Saltatory conduction along preliminarily- myelinated sensory axons^45^ and conduction velocities in micropatterned (<1 µm) hSN networks^46^ were achieved in 2D, but face biophysical limitations as they do not integrate physiologically-relevant microenvironments^47^. On the 3D-scale, hydrogels patterned with microtechnology techniques were used to guide axonal growth^32,48–50^. However, despite efforts to achieve higher spatial resolution of guidance cues^51,52^, these systems are often limited by insufficient resolution of patterning features (>10 µm). Additionally, the use of microfluidic devices on the 10s-100s µm scale have been demonstrated to successfully confine and guide hydrogel-based cell cultures^29^. These approaches offer cellular organization and improved functional assessment, particularly by enhancing the signal-to-noise ratio of electrophysiological measurements by confining the electrical signals into microtunnels^53^.

Here, we describe a novel microfluidic nerve-on-chip ‘hydroMEA’ platform that addresses the described limitations by combining: (i) reliable myelinated system with human cells, (ii) precise spatial organization through microfluidic control, (iii) physiologically-relevant microenvironment, and (iv) high-resolution electrophysiological measurements for long-term studies. The described system incorporates photocrosslinkable hydrogels, which provides a tunable 3D microenvironment that mimics the mechanical properties of neural ECM, and supports a more- physiological cellular morphology and function. Co-cultures of hSCs with hSNs can be supported for over 60 days *in vitro* (DIV), and these targeted interactions promote reliable myelination. It also includes microchannels that serve as local electrical signal amplifiers, and enhance the capabilities of HD-MEA based measurements to enable reliable electrophysiological recordings from 3D-grown with unprecedented reliability and throughput. By combining biological relevance with engineering control, the hydroMEA represents a substantial advance in neural microphysiological systems. The ability to directly measure the functional impact of myelination on signal propagation paves the way for investigating myelin-related diseases and evaluating potential therapeutic interventions. Furthermore, the integration of human-derived cells enhances the translational potential of hydroMEA, and addresses a critical need in preclinical assessment of neurotherapeutics.

## 2. RESULTS AND DISCUSSION

### 2.1 A Hydrogel-based Microfluidic Platform to Facilitate Electrophysiology of Neuron and Schwann Cell Co-cultures

In this work, we introduce hydroMEA, a physiologically-relevant 3D MPS to study neuron-glia interactions and enable peripheral nerve myelination. HydroMEA integrates hydrogels, which are tuned to match the mechanical properties of neural ECM, into a microfluidic polydimethylsiloxane (PDMS) guiding structure, which can be placed either onto a glass substrate or HD-MEA. HydroMEA enables the long-term culture (*>*DIV 80) and electrophysiological analysis of myelinated hSNs (**Figure 1**). Our platform uniquely recapitulates key aspects of the *in vivo* neural microenvironment while providing <1 µm spatial control of the networks, to address limitations of existing *in vitro* models. The PDMS structure features nine identical parallel networks, enabling parallelization of experiments on a single commercial HD-MEA (**Figure 1A**), with the microstructure dimensions detailed in **Figure S1**. Each individual network incorporates a central seeding well designed to accommodate a spheroid of hSNs (**Figure 1B**). Eleven microchannels (4 µm height × 10 µm width) per side extend bilaterally from this central seeding well to guide axonal outgrowth, while assuring just few axons outgrowth per channel and good signal-to-noise ratio due to the locally-confined electrical field. The microchannels feature a 1.5 µm narrow constriction on both ends that prevents migration of hSN soma (green) into the microchannels. Microchannels then connect to lateral seeding wells, into which hSC spheroids (yellow) are seeded. At both ends, there is a wider ‘nerve-forming’ channel (30 µm height x 30 µm width) to allow hSCs migration and their controlled interactions with the extended axons of hSNs and the hSCs. This closely resembles the *in vivo* organization of myelinated peripheral nerves. When placed on top of a HD-MEA, the microtunnels are aligned with underlying electrodes and ensure that each action potential is acquired (**Figure 1C).** The hydrogel (blue) is confined inside the microtunnels and seeding wells. After seeding spheroids in the seeding wells, the cells grow their axons into the microtunnels while the hSN cell bodies remain in place, offering confined yet 3D growth (**Figure 1D, 1E).** Interestingly, while the neuronal cell bodies were unable to enter the microchannels, the hSC were able to squeeze through the small openings and migrate throughout the microstructure.

**Figure 1:**
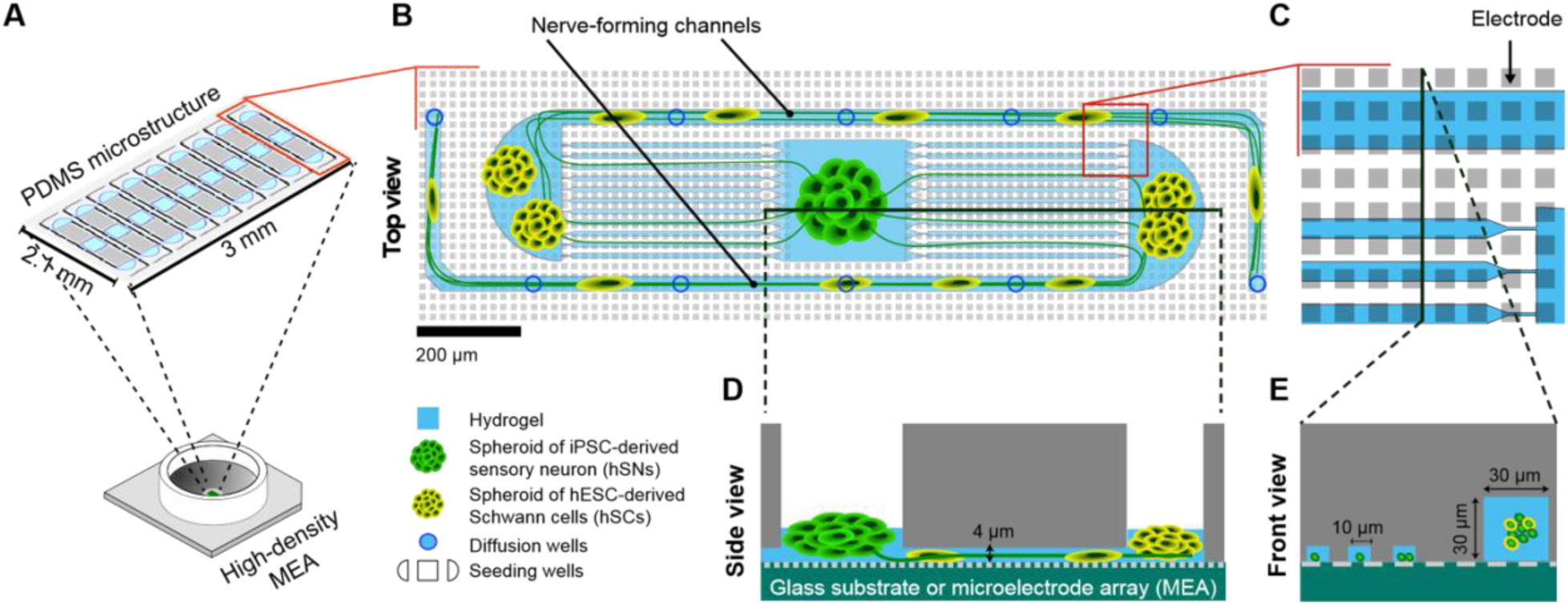
**Schematic of hydroMEA, a 3D hydrogel-based myelinated-nerve-on-chip platform**. **A**: The PDMS microstructure features 9 parallel identical networks that fit the sensing area of the MEA array. **B**: Zoomed-in top view of one network. The design comprises a central seeding well to host a hSN spheroid, from which eleven 4 µm×10 µm microchannels project from each side onto lateral seeding wells hosting hSC spheroids and each leading to one single 30 µm×30 µm nerve-forming channel. **C:** Zoomed-in view to show microchannels some of which narrow down to 1.5 µm in order to limit hSCs migration. **D:** Cross section of one side of the network. hSCs migrate out of the spheroid into microchannels and the nerve-forming channel and interact with the axons. The whole network is densely covered by microelectrodes (grey squares) that can be used to stimulate or record electrical activity. **E:** Cross section of **C** along the indicated line. The nerve-forming channel is higher and wider than the microchannels. The height of the last segment of the nerve-forming channel decreases down to 4 µm to enhance the axon bundle stimulation.

### 2.2 GelMA-based Hydrogels Match the Mechanical Properties of Neuronal ECM

Bulk hydrogels of various chemical compositions were screened to evaluate materials that could match the mechanical properties and structure of neural tissues. Three compositions of hydrogels, including (i) alginate, a polysaccharide for its independent tuning of mechanical properties; (ii) gelatin methacryloyl (GelMA); and (iii) polyethylene glycol (PEG) with matrix metalloproteinase (MMP)-cleavable peptides (PEG-MMP), were prepared, tested, and compared on how they supported growth of hSNs for >2 months (**Figures S2, S3, S4)**. Either the percent weight/volume (w/v) of each hydrogel precursor, or the amount of crosslinker were independently tuned to modulate the resulting mechanical properties. While alginate was crosslinked in the presence of a divalent cation (e.g., Ca^2+^), GelMA and PEG-MMP were crosslinked with a photoinitiator (LAP) and exposure to near-UV light (405nm). The exposure time for all the GelMA and PEG–MMP formulations was kept constant (100 seconds). Alginate formulations of higher molecular weight (HMW, more elastic) and lower molecular weight (LMW, more viscoelastic) were compared, as well as formulations that were first freeze-dried before crosslinked to create macroporous structures, as previously described^26,54^. After their reliable fabrication was established, the hydrogels were seeded with random cultures of a primary rat cortical neuron suspension and evaluated for neurite outgrowth. The LMW alginates were able to best support neurite outgrowth at DIV 13, with increased neurite coverage on the microporous formulations (**Figure S3**). For the photocrosslinkable hydrogels, a 5% w/v formulation of GelMA was found to best support neuronal growth, particularly with increased concentrations of laminin mixed into the precursor solution. While the viability of neurons in PEG–MMP hydrogels was adequate at DIV 14, there was considerable loss of neurites at later timepoints (e.g., DIV 48) which indicated a lack of sustained neural cell support with this composition. These hydrogels exhibited poor neuronal outgrowth and strong neuron body clustering (**Figures S2, S3**), and thus only alginate and GelMA were considered for further studies.

While both the alginate and the GelMA offered sufficient neuronal support, an additional material requirement for the proposed platform was the ability to process the formulations for several minutes to sufficiently fill the small 1 µm compartments. The alginate formulations crosslinked almost immediately (<15 seconds) in the presence of the divalent cation, while the GelMA precursor solution could fill any desired shape and then be crosslinked into a hydrogel in the presence of near-UV light. As a result of the increased working time, the GelMA formulation was selected to grow neurons in bulk and micropatterned hydrogels.

As the presence of laminin demonstrated improved viability and neurite outgrowth, GelMA formulations were supplemented with a consistent amount of laminin (0.11 µg/ml). After exposure to 405nm light for 100 seconds (**Figure 2A**), transparent hydrogels were formed (**Figure 2B**). To evaluate the optimal GelMA formulations for the hSNs, compositions of 2.75–10% w/v GelMA were fabricated. However, the 2.75% w/v GelMA formulations were extremely fragile and did not provide sufficient mechanical integrity for mechanical testing on a rheometer. As a result, formulations between 3.25–10% w/v GelMA were measured (**Figure 2C**). At least 3 hydrogels of each formulation from one batch, with at least 3 independent batches, were fabricated and measured to quantify the storage (G^′^) (**Figure 2C(i))** and loss (G^′′^) (**Figure 2C(ii))** moduli. As the w/v % of GelMA increased, the hydrogel G^′^ increased. A similar trend was observed for the G^′′^, though there was increased variability in the lower % w/v formulations. To minimize batch variability, the precursor solution was mixed every 2 minutes with a pipette set to at least half the volume of the precursor. The precursor was also kept on ice until the entire volume was used, to minimize reactivity of the photoinitiator. As GelMA hydrogels between 5-6% w/v were found to match the range of healthy nerve tissue (G^′^∼300 Pa^55^), a 5% w/v GelMA composition supplemented with laminin was selected to culture the hSNs.

**Figure 2:**
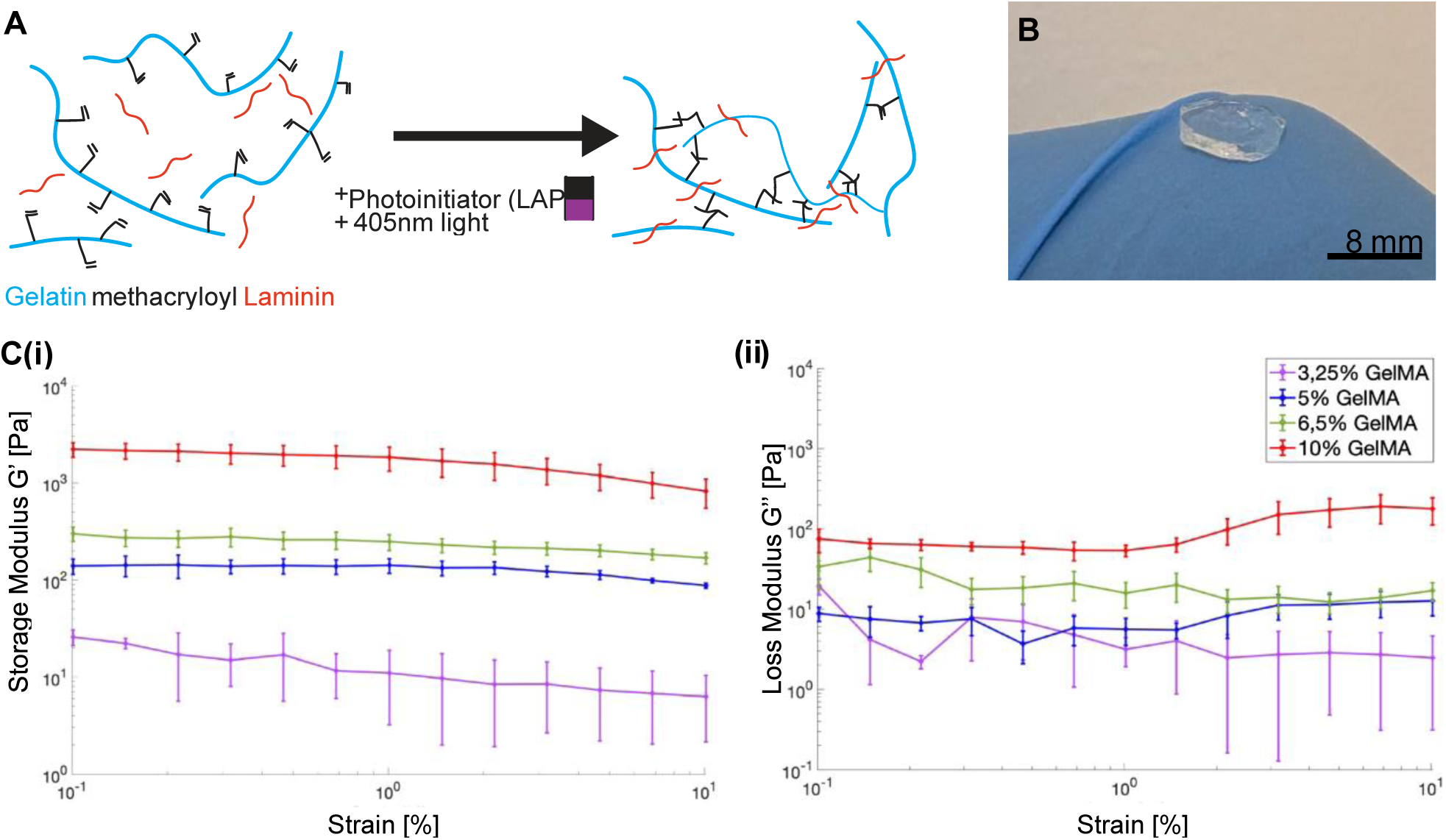
Fabrication and mechanical characterization of GelMA across various % w/v. **A**: Laminin can be incorporated into the gel during the photocrosslinking process. **B**: Photograph of the final gel material. **C**: Strain-dependent storage (i) and loss modulus (ii) of various GelMA formulations. Each formulation has n=9-11 different hydrogels, with at least 3 separate batches. Mean and s.d. are plotted.

### 2.3 Optimized GelMA-based Scaffolds Maintain Viable Co-cultures

To investigate the suitability of the optimized GelMA formulations for *in vitro* neural applications, the hydrogel was first used to support long-term bulk 3D culture of hSNs, hSCs, and hSNs-hSCs co-cultures. The fabricated GelMA hydrogels successfully supported the long-term (>DIV 70) viability and growth of hSNs (**Figure 3**). Spheroids of hSNs demonstrated robust survival and extensive axonal outgrowth when cultured in bulk GelMA hydrogels for 30 days and stained for a live-cell marker, calcein AM (**Figure 3A(i)-(iii)**). At DIV 60, hSNs had extended dense neurite bundles into the entire volume of the hydrogel matrix, as visualized by immunohistochemistry (**Figure 3C**). Additionally, hSCs demonstrated similar compatibility with the GelMA hydrogel formulation (**Figure 3B(i)-(ii)**). In contrast to hSNs, hSC morphology and behavior appeared relatively indifferent to the stiffness of the bulk hydrogel across the tested range (**Figure S7**). More importantly, both hSNs and hSCs maintained viability and appropriate morphological characteristics >DIV 30, which established the effectiveness of the proposed hydrogel. Further, the hydrogel could support long-term co-cultures which was important for the investigation of mechanistic phenomena such as myelination, which occur over prolonged periods (>6 weeks).

**Figure 3:**
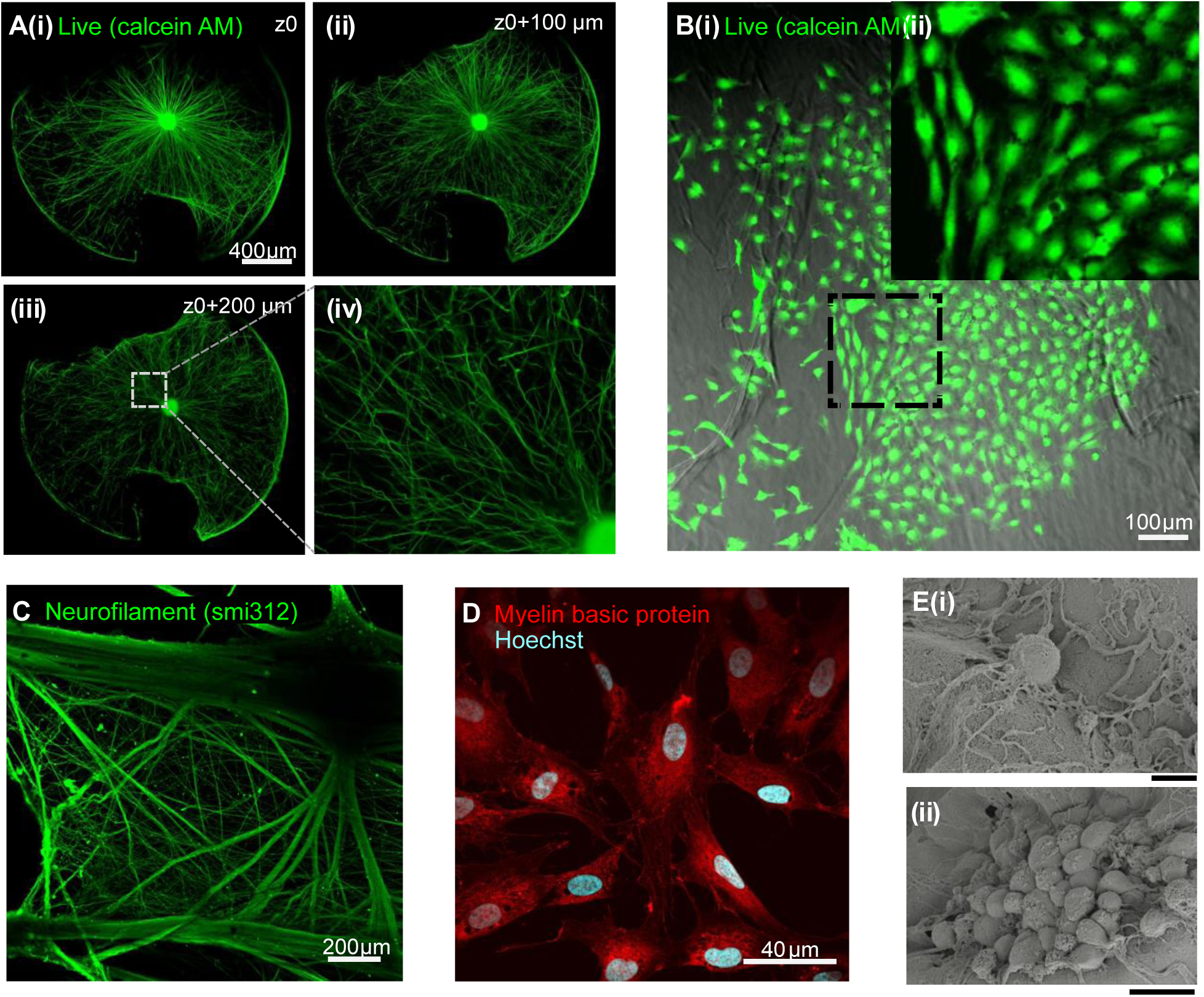
**Optimized GelMA-based bulk gel supports 3D growth and viability of hSNs and hSCs for long-term culture. A:**(i)-(iii): 3D culture of a spheroid of hSNs in a GelMA-based bulk gel (DIV 30) for three different z planes of the gel, including the (i) neutral plane, (ii) +100µm in z, and (iii) +200 µm in z. (iv): Zoomed-in view showing the 3D tortuous growth of axons in the gel, in a single plane. Cells are stained with calcein AM (green) to visualize the live cells. Scale bar = 400 µm. **B**(i): hSCs are cultured on a GelMA- base bulk gel at DIV 9. Scale bar = 100 µm. **C**: Immunostained 3D grown hSNs in bulk gel (DIV 55). Cells were stained for neurofilament (SMI312, green). Scale bar = 200µm. **D**: Immunostained hSCs grown in a 3D bulk gel at DIV 60. Cells were stained for myelin basic protein (MBP, red). Scale bar = 40 µm. **E**(i-ii): Scanning electron microscopy photomicrographs of (i) a single hSN cell body, with the scale bar = 5 µm and (ii) a spheroid of hSNs, both in GelMA-based bulk gel, with the scale bar = 20 µm.

The stiffness of the GelMA hydrogel influenced both axonal growth characteristics and bundling behavior (**Figure S4**), as stiffer gels led to decreased neurite outgrowth and bundled growth. A particularly striking observation was the distinct morphological differences between hSNs cultured in 3D versus on top of conventional 2D, rigid glass substrates. Axons grown in the hydrogels exhibited a meandered growth pattern, where the axon moved in and out of different planes and sharply contrasted with the linear trajectories observed on glass substrates (**Figure 3A(iv), Figure S5**). This observation was also confirmed by scanning electron microscopy (SEM) (**Figure S6**). This tortuous growth pattern was consistent with previous studies which demonstrated that 3D environments more accurately recapitulated the complex spatial constraints encountered by axons *in vivo*^56,57^. Neuronal cell body morphology was also influenced by the mechanical microenvironment. When grown in the 3D hydrogel, the hSN soma had a rounder, enlarged morphology compared to the flattened phenotype of hSN observed on 2D cultures (**Figure 3E**). This change in cellular morphology would affect neuronal mechanobiology and function^58^. Further, the 3D morphology and phenotype closely resembled the morphology of *in vivo* chick embryo sensory neurons at day 18 of development, as previously reported^59^, which confirmed the importance of a physiologically-relevant mechanical cues for maintained native cellular behaviors.

To further explore the effects of the mechanical environment, neuronal cultures were evaluated with two distinct approaches: (i) hSNs seeded on top of the formed hydrogels, and (ii) hSNs encapsulated within the hydrogel precursor, to be crosslinked into the hydrogel matrix. While full encapsulation appeared to delay the initial onset of neurite extension compared to surface seeding, hSNs demonstrated a remarkable ability to rearrange and infiltrate the hydrogel matrix during subsequent growth phases when seeded on top of the formed hydrogels (**Figure S5**). This behavior indicated that the neurons actively remodeled their microenvironment, consistent with the dynamic characteristics of cell–matrix interactions observed during neural development.

To investigate the functional interactions between hSNs and hSCs, co-cultures of the two cell types were grown within the hydrogel (**Figure 4**). The GelMA-laminin hydrogel not only supported individual cell type viability and growth, but also facilitated the complex physical interactions necessary for proper neural function. hSCs successfully integrated with the neuronal networks, and a few aligned to the extended axons in both random mesh-like arrangements (**Figure 4A**) while many more hSCs were aligned to the organized bundle-like axonal structures (**Figure 4B, 4C**). Importantly, hSCs exhibited active interaction with axons, demonstrated characteristic alignment along axonal trajectories, and in several instances, wrapped around isolated single axons and axonal bundles (**Figure 4C, 4D, Figure S8**). After DIV 60, immunohistochemistry assessment of the co-cultures revealed myelinated axons within the bulk hydrogels (**Figure 4E**). Interestingly, this myelination occurred even though the axonal diameters were occasionally <1 µm, a threshold previously considered for peripheral myelination^60^. This could be because the 3D hydrogel mechanical properties offered an appropriate microenvironment to promote and facilitate myelination through mechanisms beyond diameter-dependent selection, and rather through enhanced cell–cell signaling or altered mechanical constraints that promoted hSC differentiation and myelination.

**Figure 4:**
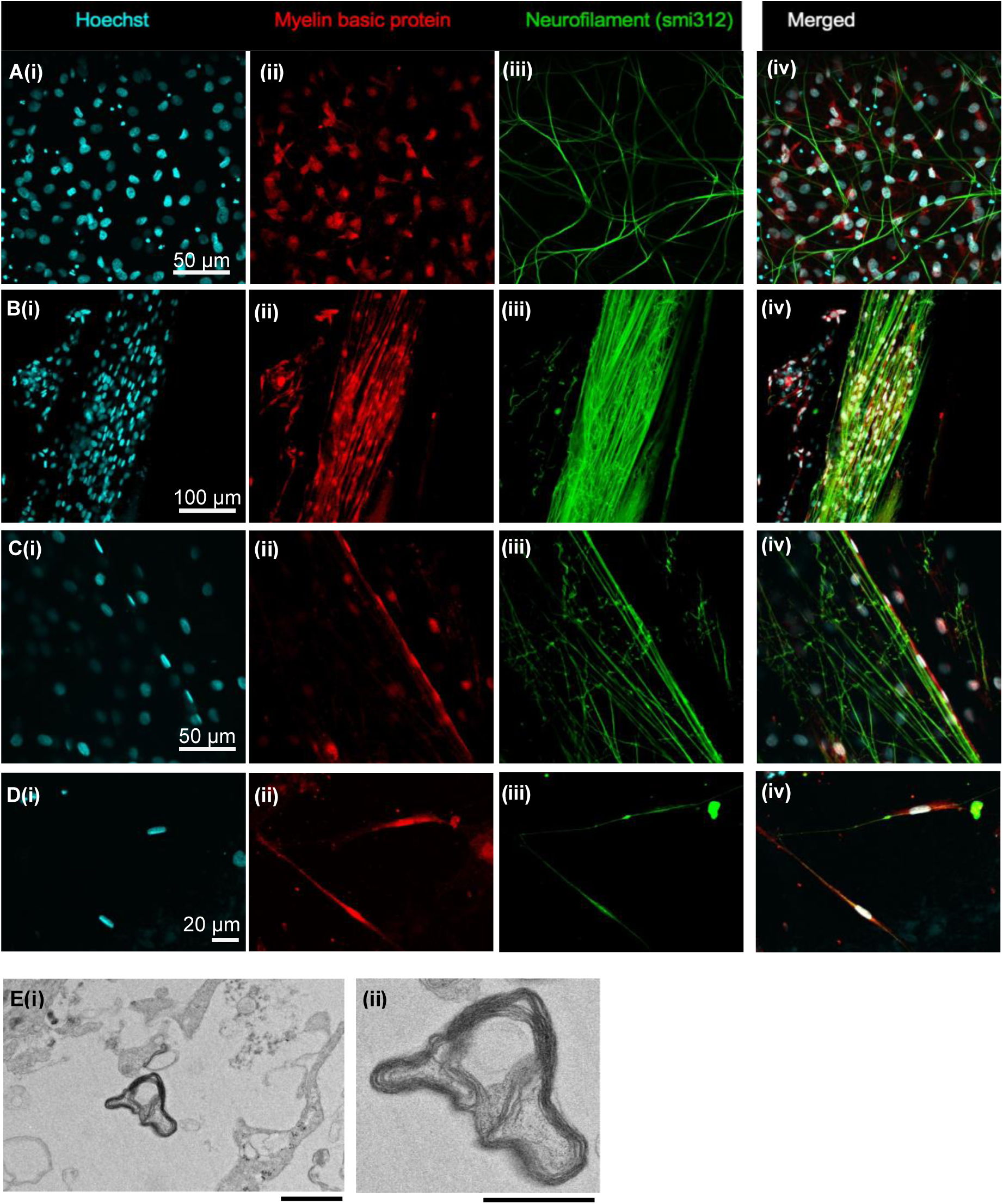
**GelMA-based bulk gel support 3D grown and viable co-culture of hSNs and hSCs while promoting co-culture interactions. A-D**: Images of stained co-cultured hSNs and hSCs in GelMA-based bulk hydrogel. Different cellular organizations are shown. **(A):** Unorganized mesh-looking co-culture. Scale bar = 50 µm. (**B):** hSCs infiltrated inside a bundle of axons. Scale bar = 100 µm. **(C):** hSCs elongated and wrapped long a few bundled axons. Scale bar = 50 µm. **(D):** Single hSCs elongated and wrapped along single isolated axons. Scale bar = 20 µm. Cells were stained for Hoechst (cyan), myelin basic protein (MBP) (red), neurofilament (SMI312) (green). All samples were fixed at DIV 56. **E**: (i) Transmission electron microscopy image of a myelinated axon taken at DIV 74. Scale bar = 1 µm. (ii) Zoomed-in view of the myelin rings. Scale bar = 500 nm.

### 2.4 3D Hydrogel-Integrated Microstructures Enable the Evaluation of Neuron-Schwann Cell Interactions

After the identification and determination of the bulk hydrogel conditions to support hSNs and hSCs, the next goal was to enable a precise spatial network. This was achieved with a microfabricated PDMS microstructure, which was then filled with the described hydrogel formulation (**Figure 5A**). As the PDMS was hydrophobic, the surface had to be made hydrophilic through oxygen plasma activation. Initially, Method 1 explored the plasma treatment of both the PDMS and the underlying substrate followed by assembly of the two components, but this method did not provide sufficient bond stability over time (**Figure S9**). However, in Method 2, when the microstructures were first adhered onto the underlying substrates and then the pre-assembled construct was exposed to oxygen plasma, the maintained van der Waals-based bonding promoted reliable adhesion over multiple weeks (**Figure S10**). As a result, Method 2 facilitated the uniform filling of the microchannels with the GelMA precursor, which was then photocrosslinked *in situ* after completely filling the microstructured PDMS (**Figure S11**). When compared to 2D cultures which had the same microstructured PDMS but were instead coated with a thin layer of a silk-based coating (iMatrix^TM^) to promote cell attachment (**Figure 5B**), the 3D hydrogel-filled constructs exhibited an altered axonal outgrowth (**Figure 5C**). In both cases, the axons of hSN spheroids had extended into the entire microstructure by DIV 25 while the neuronal soma remained confined in the seeding compartment. While the axons of hSNs cultured on 2D-coated glass surfaces grew individually across the glass surface with considerable varicosities (**Figure 5B**), the hSNs cultured within the 3D hydrogel environment exhibited pronounced axonal bundling and fascicle formation, as they grew inside the hydrogel (**Figure 5C**). This suggested that the hydrogel properties modulated the biomechanical forces that govern fasciculation^61^. Importantly, when hSCs were added to their defined compartment, the hydrogel-filled microchannels enhanced hSC-axon interactions as the hSCs migrated into the nerve-forming channels by DIV 12 (**Figure S12**). Moreover, the hSCs grown in 3D contacted, aligned, and began to wrap around axons, which was a sign of in early myelination.

**Figure 5:**
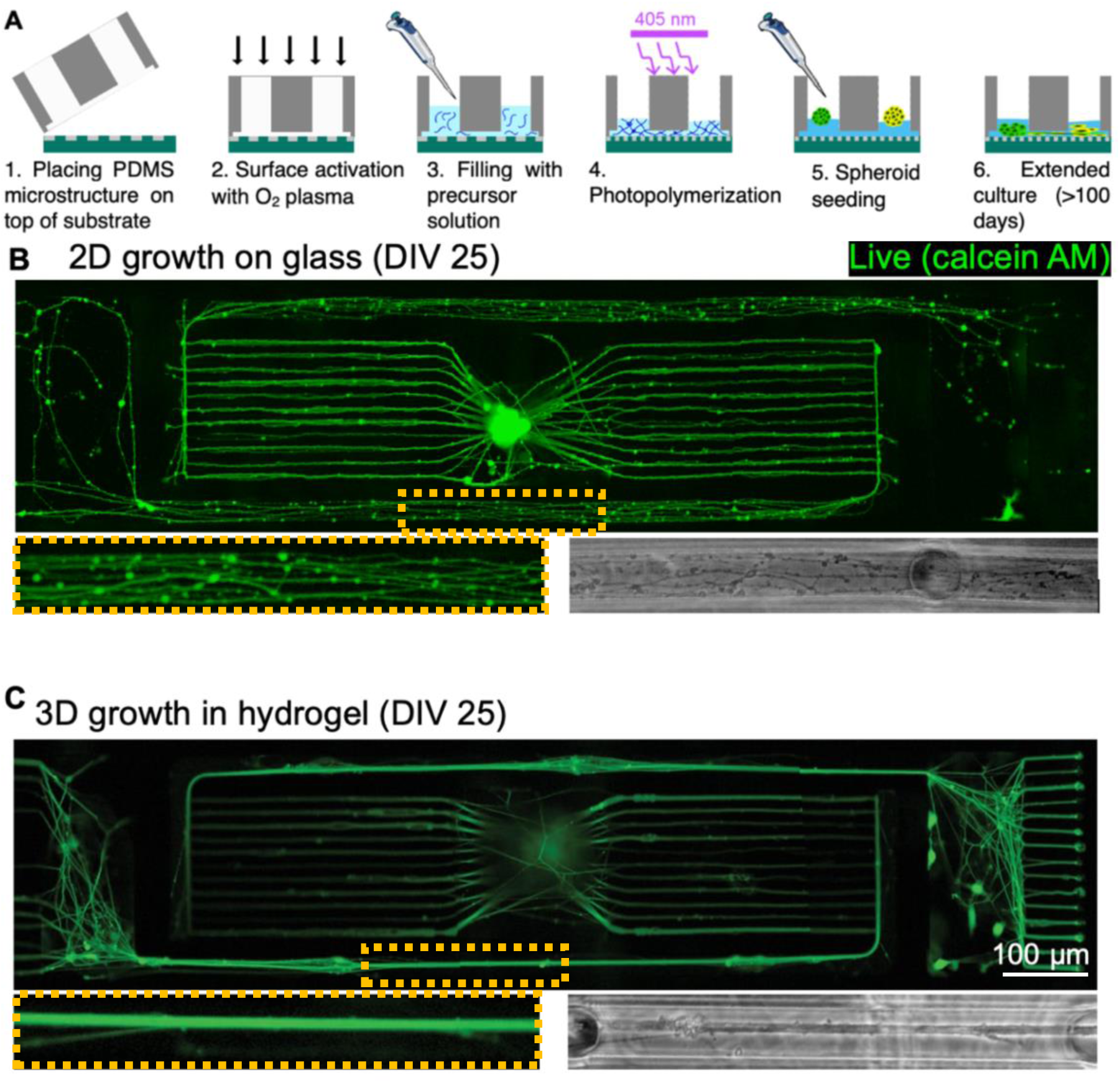
GelMA-based hydrogel integrated in the PDMS-microstructure enables 3D guided growth of hSNs. **A**: Schematic depicting the experimental process flow to integrate hydrogel inside PDMS microstructures on a glass substrate or on a microelectrode array (MEA). **B**: A fluorescence image of hSNs grown in 2D on iMatrix-511 silk-coated glass substrate in the guiding microstructures is shown on top. A fluorescence and a phase contrast zoomed image of the highlighted rectangular area (orange) are shown below. **C**: The comparative images when hSNs were growing in the 3D hydrogel matrix show axons bundled together. Images in **5B** and **5C** were taken at DIV 25 and stained for live-cell dye, calcein AM (green).

To better characterize these complex co-culture interactions, immunohistochemistry was used. Axons were stained with peripherin, to visualize unmyelinated axons (**Figure 6, Figure S13**). While some axons were peripherin-positive, the axons that were wrapped by hSCs and visualized by myelin basic protein (MBP), were absent of the peripherin stain which suggested active selection of specific axon populations for myelination. This was consistent with previous reports, which showed that only a subset of axons undergo myelination *in vivo*^4,60^. The observation also demonstrated the unique capability of the described hydroMEA platform, to combine human-derived cells, physiologically-mimicked 3D microenvironments, and geometric confinement to facilitate imaging of defined regions for co-culture interactions and myelination.

**Figure 6:**
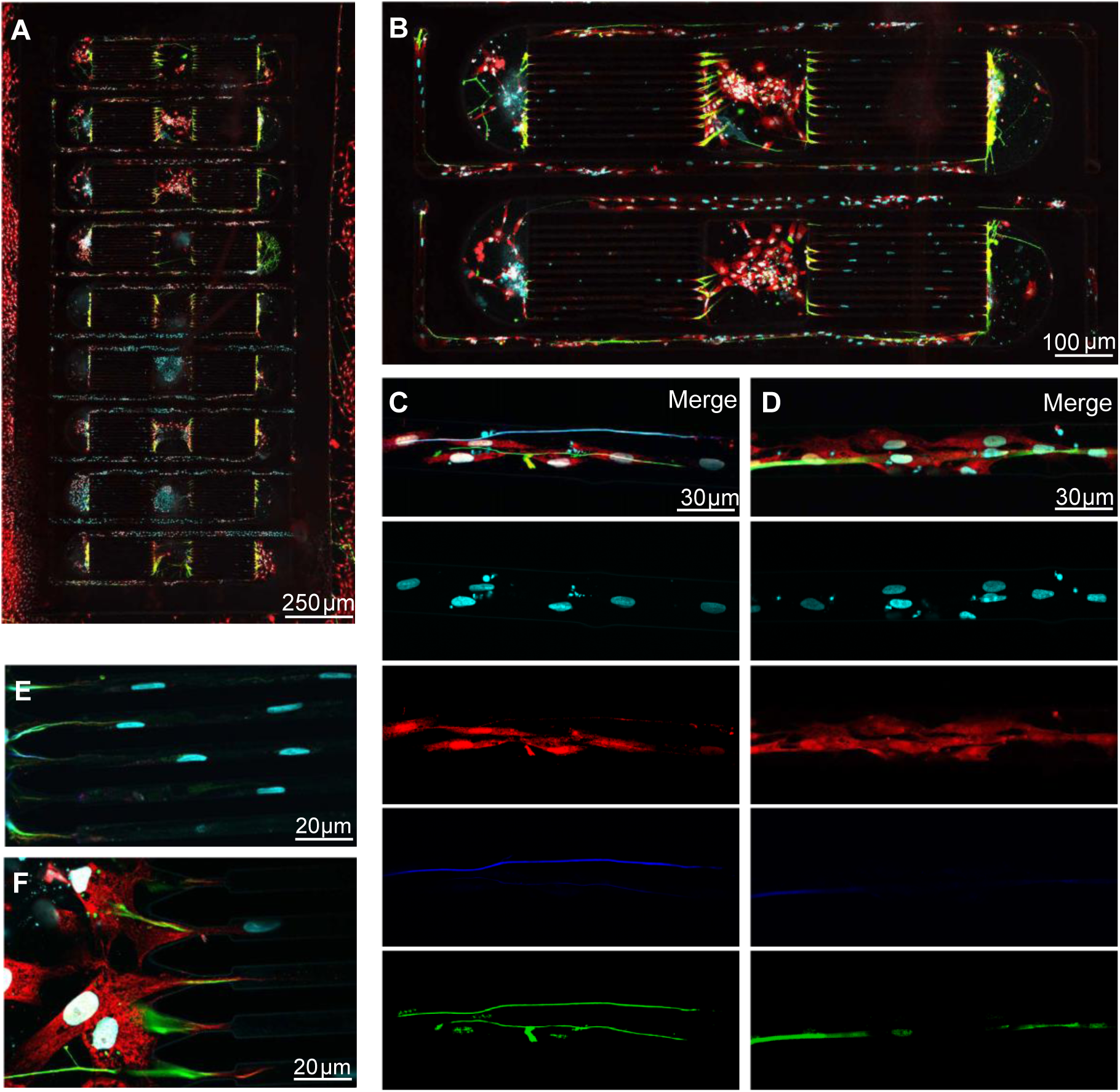
Co-culture of hSNs and SCs can be grown in 3D and in a guided fashion in hydrogel-integrated PDMS microstructure at DIV 56. **A**: Image of nine parallel networks featuring hSNs and hSCs in microstructures. Scale bar = 250 µm. **B**: Zoomed-in view of two networks from **A**. Scale bar = 100 µm. **C**: Zoomed-in view of B of one nerve compartment portion exhibiting selective alignment of SCs along an axon. Scale bar = 30 µm. **D**: Zoomed-in view of **B** exhibiting alignment and wrapping behavior of SCs around axons. Scale bar = 30 µm. **E**: hSC nuclei shown in cyan (Hoechst) deform, adapting to the narrow microchannels. Scale bar = 20 µm. **F**: hSCs start entering microchannels. Scale bar = 20 µm. For all subpanels, cells are stained for neurofilament (green), peripherin (dark blue), MBP (red), and Hoechst (cyan).

### 2.5 Electrophysiological Assessment of Schwann Cell Mediated Axonal Conduction Properties

Finally, in addition to the qualitative structural validation of myelinated axonal segments with immunohistochemistry staining and imaging, the quantitative functional effects of the SCs were evaluated by measuring the conduction velocities of the axons on the hydroMEA (**Figure 7**). Hydrogel-filled microstructure constructs were prepared as described above, but the microstructured PDMS was instead placed on top of a HD-MEA rather than on a glass substrate. The hSNs and SCs were seeded in their designated compartments, and starting at DIV18, the underlying electrode at the far-end of the axon bundles was used to electrically stimulate the axonal ends (**Figure 7A**). This external stimulation then elicited action potentials in the recruited axons, which could be measured in the designated recording microchannels. As the microchannel length (distance) was defined by the microstructure geometry, and the time difference between the stimulation and arrival of the elicited action potentials was recorded, the conduction speed of the propagated action potentials could be inferred from signal latency. Since the stimulation was delivered as a train of repeated pulses, other activity-dependent metrics such as conduction slowing and failure behavior could also be reliably measured^46^ (**Figure 7B).** At the stimulation site, 1000 pulses were delivered and 1000 pulse-induced spikes were recorded at each frequency (**Figure 7B(i)).** There were two distinct bands present (**Figure 7B(ii), Figure 7B(iv))** which suggested 2 axons were recruited in the specific microchannel shown.

**Figure 7:**
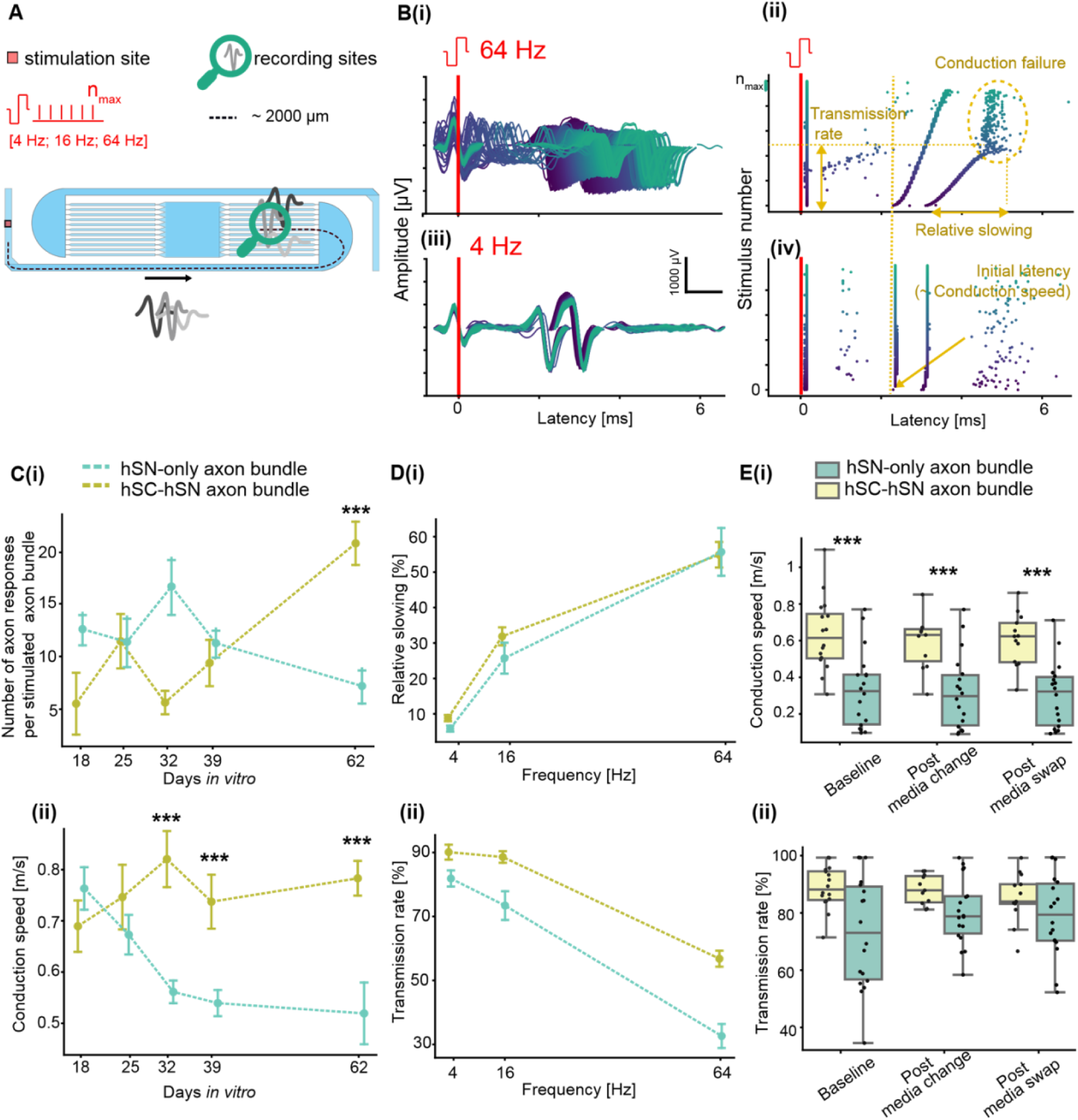
Electrophysiological characterization of myelinated nerve-on-a-chip. **A**: Stimulation protocol to measure stimulation-induced nerve responses. Each nerve end is stimulated by a train of n pulses at different frequencies, and elicited action potentials travel down fibers and are measured in recording microchannels. **B**: Post-stimulus raster plots showing example responses in one single microchannel upon nerve stimulation at 4 Hz (bottom) and at 64 Hz (top). These post-stimuli raster plots are the basis of electrophysiology metrics extraction. **C**: (i) Number of responding axons per nerve over days in *vitro* and (ii) Conduction speed per nerve over DIV 64. **D**: (i) Relative response slowing over increasing frequencies. (ii) Transmission rate over increasing frequencies. **E**: (i) Conduction speeds across different media change intervention and (ii) Transmission rate across different media change intervention. Each data point is the average of individual axon responses per stimulated bundle. In all quantifications, at least n = 16 bundles. All data are shown as mean ± s.e. ***: p*<*0.001 (Welch’s t-test).

At DIV 62, the number of responsive axons that were stimulated in the hSN-only axon bundle slightly decreased, while the axon bundles in the hSN-hSC showed increased axonal responses, and reached an average of 20 responses per axon bundle at DIV 62 (**Figure 7C(i), Figure S15A**). An additional experiment with hSN-only axon bundles was monitored over 99 days, and partially confirmed this trend although (**Figure S14**). While the sample preparation and spheroid seeding protocols were highly reproducible, individual networks and MEAs exhibited variability in the rate of axonal growth, behavior.

Conduction velocity (*ν_c_*) is a critical parameter to assess axonal conduction and is expected to be faster in the presence of Schwann cells and functional myelin. This parameter has rarely been measured in neuron-Schwann cell co-cultures, and only with low throughput^45^. While both the hSN-only bundle and the hSN–hSC bundles exhibited similar averaged conduction speeds at DIV 18, the hSN-only cultures showed decreased *ν_c_* over DIV 64 (average *ν_c_* = 0.5 m/s) while the hSN–hSC cultures demonstrated significantly higher *ν_c_* over DIV 64 (average *ν_c_* = 0.8 m/s) (**Figure 7C(ii), Figure S15**).

The activity-dependent properties were further analyzed to gain additional insights into axonal function by evaluation of axonal fatigue. Although relative *ν_c_* slowing showed no significant differences between conditions (**Figure 7D(i)**), the hSN–hSC axon bundles exhibited reduced conduction failure compared with hSN-only axon bundles when stimulated at 16 or 64 Hz (**Figure 7D(ii)**), which suggested that the hSCs sustained the axonal response to repeated activation during high-frequency activity. This measure of transmission rate was intrinsically connected to axon excitability, and the result was in accordance with recent findings that demonstrated that Schwann cells promote neuronal excitability through enhanced sodium channel expression^62^. This hypothesis was further supported by increased spontaneous activity in only the hSN–hSC axon bundles, which indicated increased transient excitability (**Figure S16**).

To determine whether these observed differences were attributed to soluble factors in the culture medium rather than direct cell–cell interactions, the same stimulation protocols were performed and then the media from the two samples was exchanged. Briefly, the media from all the samples was collected and stored, and then replaced with fresh medium of the same relevant composition for the sample at DIV 62. After the stimulation assay was performed on all networks for 1 hour, the original media for each hydroMEA was added to the other hydroMEA; specifically, the original hSN-only media was placed in the hSN–hSC hydroMEA, and the original hSN–hSC co-culture media was added to the hSN-only hydroMEA. This allowed the characterization of whether the conditioned medium of the co-cultures, and thus relevant soluble factors could explain the measured differences in the recorded conduction velocities. Interestingly, the previously-measured *ν_c_* and relevant differences between the two conditions persisted throughout these media exchanges (**Figure 7E(i)**), which indicated that the difference in axonal conduction velocities were not primarily mediated by soluble factors secreted by the hSCs into the medium. Further, the conduction failure rates for both conditions converged toward similar values following media exchange, particularly in the swapped media condition (**Figure 7E(ii)**), which suggested that while some aspects of axonal conduction may be influenced by soluble factors, the fundamental Schwann cell enhancement of conduction velocity in co-cultures resulted from direct cellular interactions, or non-diffusible modifications to the axon microenvironment. The different media change interventions did not have any effect on the other conduction metrics (**Figure S15**). These findings aligned with a previous study^63^ which demonstrated that hSCs led to enhanced axonal conduction properties from DIV 17 even prior to the formation and visualization of myelin, but not in hSN-only bundles. The significantly faster *ν_c_* in the hydroMEA hSN-hSC co-cultures by DIV 62, as compared to DIV 18, was likely from the myelination of the axons. Thus, hydroMEA could be used to further investigate these effects and mechanisms of myelination or demyelination on *ν_c_*.

## 3. Conclusions

Overall, the combination of the optimized GelMA hydrogel with precisely controlled microfluidic architectures provided both biological relevance and engineered control over hSNs-hSCs co-cultures and enhanced the translational potential of the platform. First, the observed morphological similarities of hSNs and hSCs to *in vivo* cells, particularly the 3D cellular morphology, organization, and cell–cell interactions, suggested that the hydroMEA provided more translatable insights compared to other *in vitro* platforms. Second, the microchannels served the dual purpose of guided cellular organization and enhanced electrophysiological signal detection, a critical capability for functional assessment and monitoring of neural activity. The co-culture of hSNs and hSCs supported long-term maintenance (>DIV 100) of viable, functionally-active neural networks, and enabled myelination, confirmed with both qualitative imaging techniques and quantitative, functional evaluation through electrophysiological measurements. These results provided a comprehensive qualitative and quantitative framework to assess axonal functionality, and could be used in future studies to further evaluate the effects of myelination in neuronal cultures. Further, in addition to mimicked developmental conditions, the hydroMEA platform could be used to study diseased states, such as multiple sclerosis and other demyelination disorders, to understand both mechanisms and effects of various neurodegenerative disorders. Additionally, the hydroMEA could be used to evaluate potential therapeutic interventions, with the capacity to detect subtle but functionally significant changes in conduction properties that may precede or exist independently of structural modifications visible through conventional histological techniques. These could include electrophysiological effects induced by diseased human-derived cells, or in the presence of drugs or toxins, which establishes the hydroMEA platform as a powerful tool to close the gap towards the clinic.

## 4. Experimental Section: Materials and Methods

***Synthesis of GelMA***: The synthesis, purification, and preparation of gelatin methacryloyl (GelMA) followed previously published protocols^64^. Briefly, gelatin from porcine skin (type A, gel strength 300 bloom, Sigma-Aldrich G1890-500G), was added to a round bottom flask and diluted to a 10% weight/volume solution in deionized water. The solution was allowed to completely dissolve, before heated to 50℃until the gelatin was completely dissolved. Methacrylic anhydride (Sigma-Aldrich 276685) was added to the flask, at 3.89 mmol per gram of gelatin. The solution was left to spin for 1.5 hours, after which the solution was transferred to 50 mL conical tubes. The unreacted methacrylic anhydride and methacrylic acid were removed by centrifugation at 4500 g for 5 minutes. Next, the solution was decanted and diluted 2x. The solution was transferred to dialysis tubing (6-8 kDa MWCO, Spectrapor 7; Spectrum Laboratories 132665), and kept in a bath of deionized water for 5 days. The dialysis water was changed at least 2x per day. After dialysis, the pH was adjusted to the dialyzed GelMA solution to 7.4, using 1M NaHCO_3_. The solution was sterile-filtered using a 0.2 µm vacuum filtration unit (PES membrane), frozen at - 20℃, and transferred to a lyophilizer to completely dry. The dried material was characterized using a ninhydrin assay to determine the degree of functionalization. Duplicate samples of the dried GelMA were dissolved in UltraPure distilled water, heated, and then a standard curve of dilutions (*e.g.*, 1:1, 1:4, 1:9) were prepared with UltraPure distilled water. After a 2.5 mg/ml solution of ninhydrin was prepared, the ninhydrin solution was added to the GelMA and two blanks of UltraPure distilled water were included as a control. The tubes were incubated for 12 minutes at 50 ℃, and then transferred in triplicate to a plate reader to measure the absorbance at 570 nm.

***Preparation of GelMA Precursor Solution***: Initial experiments were done with commercially available gelatin methacryloyl, GelMA, (Sigma Aldrich, 900496) with 300 g Bloom strength and an 80 % degree of substitution . Approximately 100–200 mg of the dry powder was weighed out into a 5 mL Eppendorf tube, and then sterile phosphate buffered saline (PBS, Sigma-Aldrich, 10010015) was added to prepare a 10% weight/volume (w/v) stock solution of GelMA. When using the synthesized GelMA, PBS was added to prepare a 3.75 % w/v stock solution. The Eppendorf was placed in a dry-bead bath at 37 ℃ for 30–45 minutes, and the tube was gently rocked every 10–15 minutes to facilitate the dissolution in PBS. After formation of a uniform solution, the formulation was filtered with a Millex PVDF syringe filter, 0.45 mm pore size (Millipore, SLHVR334B), into an autoclaved 1.6 mL Eppendorf tube. The formulation was stored in a 4℃ fridge until ready for use, at which point the tube was moved to the dry bath (37 ℃) to allow the GelMA to warm for easier handling. Precursor solutions were prepared by mixing GelMA stock solution, PBS, recombinant Laminin-511 E8 fragment (iMatrix-511 Silk, Anatomic, AMS.892 012), and a photoinitiator, lithium phenyl-2,4,6-trimethylbenzoylphosphinate (LAP; at intended final concentration of 0.1 wt%). LAP was synthesized as previously described^65^. The total volume of the precursor solution was kept to 50 µL and the amount of laminin (2.75 µL) and LAP (10 µL) was kept constant across all the formulations. The precursor solution was carefully homogenized by pipetting up and down with a P100 µL without creating bubbles. Similar steps were followed for the PEG-MMP hydrogels, matching previously published protocols, with the addition of RGD solution into the formulation.

***Preparation of Bulk GelMA-based Hydrogels***: Once the precursor formulations were prepared, a 10 µL droplet of the mixture was added to a glass slide pretreated with Sigmacote (Sigma Aldrich, SL2) to minimize attachment of the formed hydrogel to the glass slide. After placing two poly(methyl methacrylate) PMMA spacers of 500 µm thickness, cut 1 mm x 4 mm with a Trotec Laser Cutter, on either side of the glass slide, a second Sigmacote-treated glass slide was placed on top of the droplet, to ensure (i) uniform thickness of the hydrogel, and (ii) to minimize evaporation of the small volume and (iii) to limit oxygen exposure as oxygen inhibits the reaction. A 405 nm LED lamp (Thorlabs), calibrated to a power intensity of 20 mW*/*cm^2^, was placed on a stand directly above the glass slides. The formulations were exposed to the light for a duration of 100 s. After illumination, the top slide was carefully removed with a pair of flat-tip forceps. The forceps were then dipped into sterile PBS and used to scoop the crosslinked hydrogel from the bottom glass slide into a sterile Petri dish filled with sterile PBS. To prevent the LAP from becoming inactive due to exposure to light, the precursor solution was covered in aluminum, and a new batch of precursor was prepared every 45 minutes. All of these steps, including the crosslinking with the LED lamp, were done under a biosafety cabinet to ensure sterility of the formulations.

***Mechanical Characterization of Bulk Hydrogels***: Bulk mechanical characterization of the GelMA formulations was done with a shear rheometer (MCR 502; Anton Paar), with a Peltier stage set to 37°C. An 8 mm diameter parallel plate geometry was used for all the test, and the hydrogels prepared to be 8 mm in diameter. To avoid loading history effects, the samples underwent a preconditioning oscillatory interval (ω = 1 rad s^−1^, γ = 0.1 %, *t* = 300 s). Dynamic oscillatory strain amplitude sweeps were performed at constant frequency (1 Hz), for hydrogels that were 0.45-0.5 mm in thickness. A gap size of 0.35–0.4 mm was used for all rheological measurements, with an initial normal force ∼0.03-0.06N. A hydrated Kimwipe was placed around the plate, and underneath the hood, to provide a humid environment throughout the entire duration.

***hSN Cell Culture***: Cryovials of human iPSC-derived sensory neurons (hSNs) (RealDRG^TM^ Nociceptors, Anatomic Incorporated) and hESC-derived Schwann cells (hSCs) (from our collaborators) were removed from liquid nitrogen storage and were placed into a 37°C bath. Both cell types were thawed the same way: cells were thawed (*<*1 minute) by gently swirling the vial in the bath until there was just a small bit of ice left in the vial. After taking the cryovial into a laminar flow cabinet, the cells were gently transferred into a sterile centrifuge tube. The cryovial was rinsed with 1 mL of DMEM/F12 (Gibco^TM^, 11320033) pre-warmed to room temperature transferring its content also to the sterile centrifuge tube. 8 mL of warmed DMEM/F12 was added slowly to the cell suspension in the tube, which was then centrifuged at 300 g for 4 minutes. After the centrifugation, the supernatant was aspirated without disturbing the pellet. The cell pellet was resuspended in 1 mL of Senso-MM Maturation Media (Anatomic Incorporated, cat. 7008) to create a homogeneous cell suspension. In general, for hSN monocultures, the Senso-MM Maturation Media was kept for three days, after which the media was changed to Senso-MMx2 (Anatomic Incorporated, cat. 1032) until DIV 14, and finally changed to a custom-made medium that was composed of Neurobasal (NB) (Gibco *^TM^*) medium, to which 5 % of B27 supplement (17504-044), 1 % GlutaMAX (35050-061) and 1 % Pen-Strep (15070-063, all from ThermoFisher) were added. Growth factor supplements were freshly added to 10 mL of neurobasal medium to obtain the following final concentrations: 50 ng/mL of nerve growth factor (NGF, 450-01), 20 ng/mL of brain-derived neurotrophic factor (BDNF, 450-02), 20 ng/mL of glial-derived neutrophic factor (GDNF, 450-10), 20 ng/mL NT-3 (450-03, all from PeproTech). After performing a cell count, the cell suspension was dispensed in one or several wells of a spheroid-forming wellplate (Spherical plate 5D^®^, Kugelmeiers^®^, SP5D-24W), pre-filled with 1 mL of warm SensoMM Maturation Media, with the volume necessary to create spheroids of approximately 250 - 500 cells per spheroid. As each seeding well contained 750 microwells a total number of 250 x 750 = 187500 neurons was needed to obtain spheroids with an average size of 250 cells per spheroids. The spheroid plate was centrifuged at 50 g for 3 minutes and left in the incubator overnight. Just before seeding, PBS was replaced by Senso-MM maturation medium. Spheroids of hSNs were seeded between 12 to 48 hours after their preparation.

***hSC culture***: Human immature Schwann cells (hSC), derived from human embryonic stem cells, were generously provided by the laboratory of Lukas Sommer (Institute of Anatomy, Universität Zürich), following previously published protocols^66^, *^unpublished^*. Briefly, the cells were immediately cultured as spheroids, prepared as described above for the hSN using a Spherical plate 5D, in an expansion medium composed of Essential 6 Medium (Gibco^TM^, A1516401) and supplemented with the following expansion factors for hSCs proliferation to obtain the following final concentrations in medium: 10 ng/mL of FGF2 (human FGF-basic, 100-18B, Preprotech), 100 ng/mL of Neuregulin 1 (Human Heregulin beta1 Recombinant Protein, 100-03) and 5 *µ*M of forskolin (66575-29-9, Sigma Aldrich).

***hSN and hSC co-culture:*** With a co-culture, the Senso-MM and Senso-MMx2 media and custom media for hSNs were respectively supplemented with the hSC factors described in the previous sentence. From DIV 14 of co-culture, media had additional supplementation of ascorbic acid to reach a final concentration of 50 µg/mL (A92902, Sigma-Aldrich). Spheroids of hSNs were seeded between 12 to 48 hours after their preparation, and 24 hours after the seeding of their seeding, hSC spheroids were placed in their respective well. One spheroid was placed in the central well of each of the 9 networks in the microstructure using a 10µL pipette. In this process, additional care has to be put into not touching the microstructure, as it may cause its detachment from the microelectrode array. Culture media was changed a few hours after seeding. Half of the medium was changed twice a week.

***Design and Preparation of the PDMS Microstructure***: PDMS microstructure layouts were designed using AutoCAD (Autodesk). PDMS microstructures were fabricated by WunderliChips GmbH (Zurich, Switzerland). As shown in detail in **Figure S1**, each network features one central seeding well separated from two lateral wells by an array of eleven (per side) 10 µm-wide, 4 µm high, 430 µm-long microchannels. The channels have 3 µm-wide constrictions at the proximal end and 1.5 µm-wide at the distal end to respectively minimize the number of axons that can grow through and reduce the amount of SCs migrating towards the center. Each lateral well is connected to one 30 µm wide axon-collecting channel to allow for axons to merge and grow until a designated terminal stimulation area. The height of the end of the large channel was designed to be variable as well, with the microchannels just 4 µm tall, for increased electrical resistance and enhanced signal detected and electrical stimulation specificity by the underlying electrodes. PDMS microstructures were manually cut out of a wafer, exposed to isopropanol for 15 min, rinsed three times using sterile ultrapure water and dried at room temperature for at least one hour after aspirating as much liquid as possible. It is essential that both the microstructure and the MEA sensing area are perfectly dry for the placement described next.

***Mounting Platform with Hydrogel Integration***: The microstructures can be placed onto different substrates depending on the application. For imaging experiments, glass bottom 35 mm diameter dishes (KIT-3522 T, WillCo Wells) were used as substrate. For electrophysiological recordings of *in vitro* hSN cultures, CMOS-based HD-MEAs (MaxOne, flat surface topology with less than 0.3 µm variation, MaxWell Biosystems AG, Switzerland) were used featuring 26’400 electrodes allowing for the simultaneous recording from up to 1024 electrodes at 20kHz. Microstructures were gently held with tweezers, and carefully placed manually onto the 3.85 mm × 2.10 mm sensing area in case of the HD-MEA substrate. Before fully releasing the microstructure, the PDMS was confirmed to be in the center of the sensing area with the reference electrode surrounding the sensing area left exposed and uncovered. In order to fill the microstructure channels while overcoming the hydrophobic effect of the small PDMS features, the assembled microstructure-substrate was plasma treated to activate all exposed surfaces and turn them hydrophilic. The glass dish or MEA chip was placed with the microstructure into a plasma cleaner (Tergeo Plasma Cleaner, PIE Scientific LLC), that has been cleaned and prepared for operation immediately before this usage for more reproducibility. The microstructure was plasma activated at 5 sscm oxygen as the process gas, RF power: 25 W, pulse ratio: 50/255, for a duration of 1 minute and 15 seconds in direct mode. After bringing back the microstructure-chamber assembly in a sterile hood, the microstructure was then filled manually under a stereomicroscope for visual feedback with the prepared GelMA precursor solution. The filling was done by dispensing a drop of about 5 µL of this GelMA solution on the top of the microstructure. The microchannels can be visually checked for filling as the liquid solution travels in, as a result of the surface activation. A liquid bridge between the tip and the dispensed liquid was maintained to remove excess liquid via aspiration to prevent a thin layer of hydrogel forming on top of the microstructure that could hinder spheroid seeding later. The chamber was then immediately covered with a glass coverslip and placed under a 405 nm UV light for 100 seconds. After crosslinking, 1 mL of PBS was quickly added to the chamber to keep the hydrogel hydrated and left to wash out the leftover cytotoxic LAP for at least one hour, ideally overnight. Microstructure placement and adhesion were then assessed by collecting an impedance map of the electrode surface as previously described^67^. A few examples are shown in **Figures S9** and **S10** (Supporting Information). Briefly, a sinusoidal electrical signal (16 mV peak-to-peak voltage, 1 kHz frequency) was generated using an on-chip function generator. The sinusoidal signal was applied to the circumferential reference electrode of the MEA and the received signal at the microelectrodes was recorded. A highly attenuated signal indicated the electrodes covered by PDMS.

***Immunofluorescence Staining***: The cell-laden hydrogels, as bulk structures, and inside the microstructured PDMS, were fixed at the desired timepoint(s). First, the residual media was removed, and the constructs were rinsed once with sterile, warmed PDMS. The PBS was then removed, and enough 4% paraformaldehyde, (PFA; ThermoFisher, Catalog number 043368.9M) solution, diluted in PBS, was added to completely submerge the gels and structures. After 15 min, the PFA solution was removed and the gels were rinsed 3 times with PBS under a chemical hood. The samples were stored in PBS until further use.

To permeabilize the cells, 0.1% Triton-X (ThermoScientific, PI85111) dissolved in PBS was added to each well, dish, or MEA, and kept for 7 min. Next, six 10-minute washes with PBS were done for each sample, with the samples placed on an orbital shaker to completely wash away the residual PFA. Next, primary antibodies were added at a 1:500 ratio to a blocking buffer solution (5% goat serum, ThermoFisher, in PBS). The primary antibodies used were:

SMI312, neurofilament (mouse), Biolegend Catalog #827904 peripherin (rabbit), Abcam ab246502 myelin basic protein, MBP (chicken), Invitrogen, PA110008 myelin protein zero, MPZ (rat), Abcam ab183868 beta III tubulin, Tuj1 (rabbit), Abcam ab18207 MAP2 (chicken), Invitrogen PA110005

For each experiment, the primary antibodies were left overnight, at room temperature, on an orbital shaker set to 50 rpm. The next day, the primary solution was removed and the samples were washed with fresh PBS (at least 5 washes, spread out over a minimum of 5 hours, with 20 minutes for each wash). In some cases, the washes were left overnight.

When the washing steps were complete, the secondary antibodies (all Invitrogen): AlexaFluor Anti-Mouse 488

AlexaFluor Anti-Mouse 555

AlexaFluor Anti-Mouse 647

AlexaFluor Anti-Rabbit 488

AlexaFluor Anti-Rabbit 555

AlexaFluor Anti-Rabbit 647

AlexaFluor Anti-Chicken 488

AlexaFluor Anti-Chicken 555

AlexaFluor Anti-Chicken 647

AlexaFluor Anti-Rat 594

were added at a 1:500 to blocking buffer, placed to cover each gel, and then covered by aluminum foil. The samples were left, protected from light, on the shaker overnight. On the third day, 6 washes, each of at least 10 minutes were done with fresh PBS, followed by the addition of Hoechst 33342 (ThermoScientific, Catalog number 62249) at a 1:500 ratio to the blocking buffer solution. The Hoechst was incubated, covered by aluminum foil, at room temperature for 30-50 minutes and its presence was confirmed with the confocal before removing the residual solution. Finally, after 2 more washes with PBS, the stained gels were carefully transferred to a glass slide, covered with a drop of Prolong Gold Antifade Mountant (Invitrogen, Catalog number P36930), covered with a coverslip, and imaged with the CLSM. The hydroMEA constructs were carefully removed of all PBS, a drop of Prolong Gold Antifade Mountant was added, and a circular coverslip was placed on top.

***Microscopy Image Acquisition***: An inverted confocal laser scanning microscope (FluoView 3000, Olympus) was used to image the fluorescently labeled cultures. Fluorescence imaging of the non-fixed CMOS chips was done by removing all excess medium and using a round glass coverslip (10 mm diameter) mounted on top of the PDMS microstructure. The surface tension between the cover slip and the chip enabled us to invert the chip for imaging in an inverted microscope. For mounting in the CLSM, the CMOS chip was placed in the recording unit which in turn was mounted into a custom-made metal insert that fits into the stage of the microscope. This configuration enabled inverted mounting for imaging with a 10x objective^67^. Microscope images were processed using ImageJ. Importantly, due to their size, stained soma are brighter and thus more visible than axons on microscopy images. To enhance the intensity of the axons compared to the soma, a pixel logarithm operator was applied to all the representative fluorescent images shown in the figures of this paper. The brightness and contrast were manually adjusted to suppress background fluorescence.

***SEM microscopy***: Scanning electron microscopy (SEM) was performed using a Magellan 400 instrument (FEI Company, USA). The fixed hydrogel-cell samples were prepared for SEM, using a protocol previously developed by Tringides^32^. Briefly, after fixing the samples with PFA, they were then placed in increasing ethanol:water dilutions (e.g., 50:50, 70:30, 80:20, 90:10, 90:10, 100:0, 100:0) and left to dry overnight under a chemical hood. Next, the samples were preserved by using a hexamethyldisilazane (HMDS) (Sigma Aldrich, CAS number: 999-97-3) in ethanol, at a 1:2, 2:1, and 1:0 ratio of HMDS:ethanol. The prepared cell-hydrogels samples were mounted onto a SEM stub covered with conductive carbon tape, and were sputter-coated with a 3 nm platinum-palladium layer using a CCU-010 vacuum coating system (safematic GmbH, Switzerland). Images were acquired in secondary electron mode using an Everhart-Thornley detector at an acceleration voltage of 3 kV, spot current of 50 pA, and 0° tilt angle.

***TEM imaging***: For TEM preparation, sample processing for chemical fixation was done in a PELCO BioWave, Pro+ microwave system (Ted Pella, USA), following a microwave-assisted fixation and dehydration procedure. Fixation was done in 2% PFA and 2.5% glutaraldehyde in 0.15M cacodylate buffer (pH 7.0) with 2mM CaCl_2_ followed by post-fixation in 1% osmium tetroxide (OsO4) in the same buffer and after washing in buffer 1% Uranyl acetate (UrAc) in deionized-H2O. After washing in deionized-H2O three times, dehydration was done in a graded series of ethanol (25%, 50%, 75%, 90%. 98% and three times 100%) on ice. For embedding epoxy resin (Merck, DE), twice 25% in ethanol followed by twice 50% (total 60 min at room temperature) and 75% (30 min at room temperature and then overnight at 4 °C) was used. The steps in 100% resin (three times: 3.5, 2 and 2 hour) were performed without microwave- assistance. Polymerization was done at 60 °C for 48 hours. Ultrathin sections of 60 nm were obtained with a diamond knife (Diatome Ltd., CH) on a Leica UC6 ultramicrotome (Leica Microsystems, CH), placed on formvar and carbon coated TEM grids (Quantifoil, DE), and poststained with uranyl acetate and Reynold’s lead citrate. Ultrathin cross-sections were obtained after chemical fixation and resin embedding of the hydrogel-filled PDMS channels enveloping the cells. Stained sections were then imaged using a Morgagni 268 TEM at 100 kV in bright field mode (Thermo Fisher Scientific, USA).

***Electrophysiology set-up***: Electrophysiological recordings were performed in an incubator with a temperature set to 35℃, 90% humidity and 5% CO_2_ concentration; this setup allowed for continuous recordings up to several hours. After placing the MEA chip on the recording unit, it was allowed to rest for at least five minutes before a recording session was initiated, ensuring that any spontaneous activity disrupted by movement or changes in CO_2_ levels and temperature could return to baseline. Networks were selected utilizing the previously mentioned impedance map. A threshold was applied to identify electrodes not covered by the microstructure. A selection of about 1000 electrodes for each network from which to record electrical signals was performed using a custom-built previously published software^46^. Selected electrodes were then routed to available amplifiers and the resulting configuration was downloaded to the chip using the Python application programming interface (API) provided by the manufacturer (MaxWell Biosystems). The network activity was tracked by recording the voltage on the routed electrodes. Voltage recordings were acquired at 20 kHz sampling frequency with a resolution of 10 bits and a recording range of approximately ± 3.2 mV, which results in a least significant bit (LSB) corresponding to 6.3 µV. Using custom software based on the MaxWell Python API^67^, the raw traces were recorded and stored as HDF5-files. To apply a repetitive stimulation pattern to a network, a custom Python script was created. The script makes use of the MaxWell Python API to send commands to the system hub. A biphasic pulse with a leading cathodic phase and 400 µs pulse width was used with a stimulation amplitude of 1000 mV (2000 mV peak-to-peak). The amplitudes were rounded to the closest value available on the 10-bit DAC with approximately 3.2 mV step size.

***Electrophysiology measurements***: Electrophysiology measurements were always performed the day after last medium exchange. In the longitudinal experiment, at least 15 nerves per condition (distributed across 9 networks per chip) were stimulated and recorded at DIV 18, 25, 32, 39 and 62. The stimulation (250 pulses, 4 Hz, 1000 mV voltage amplitude), was applied first to the ends of one of the nerve-forming channels on the same networks side for each network, and then to the other networks side, so that each network does not undergo two subsequent stimulation. For the frequency ramp experiment, at least 15 nerves per condition (distributed across one chip per condition) were stimulated at DIV 62. Stimulation consisted in 1000 pulses at 4 Hz, 16 Hz and 64 Hz applied to each axon-collecting channel-end one after the other. For the medium conditioning experiment, medium was aspirated until leaving a thin film liquid above the microstructure and quickly replaced by either fresh media (in the “post media change” condition) or medium from the counter chip that had been previously removed and kept in an Eppendorf vial (“post media swap” condition).

***Electrophysiology analysis***: Analysis of stimulated-nerve responses was performed according to the analysis pipeline as previously developed^46^. Briefly, every single microchannel is physically spanned by a set of electrodes, which results in row-like arrangement of pixels on the impedance map representation of the microstructure. Therefore, microchannels were visually identified on the impedance scan and respective electrodes/pixels were manually selected via a custom-made GUI and saved in a separate file. The post-stimulus raster plot and corresponding spike waveform plot were generated for each electrode (one per microchannel) to visually inspect the response of axons in each microchannel to the various stimulation protocols. As the stimulation signal saturates the amplifiers (capped at 3.2 mV) independently from the stimulation amplitude, an artifact amplitude threshold of 3 mV was established to detect the stimulation pulse times on the stimulation electrode. Then, the best electrode for signal detection was selected based on two criteria: 1) closer to the center of the microchannel and 2) featuring a high pixel value based on the initial impedance scan (*i.e.*: the electrode is found to be more exposed to conductive medium). For each selected electrode per microchannel, spikes were detected within 0.5ms and 30ms from the stimulation artifact, for which the resulting post-stimulus raster plot and associated spike waveform can be plotted. This time window was sufficiently long for the action potentials to travel from the stimulation electrode to the microchannels. Raw data were processed using a band-pass filter (4th order acausal Butterworth filter, 300-3500 Hz). The baseline noise of the signal was characterized using the median absolute deviation for each electrode. Action potential event times were extracted from the voltage trace by identifying negative signal peaks below a threshold of 5 times the baseline noise. Successive events within 0.75 ms were discarded to avoid multiple detection of the same spike. Spike amplitudes were defined as the absolute value of the negative amplitude of the detected peak. Spike waveforms were extracted from the filtered voltage trace using the data within a −1ms to 1ms window around the timestamp of the detected spike. The waveforms were also used to measure the spike amplitude. Briefly, each induced response upon nerve stimulation was extracted via a clustering algorithm, and metrics such as number of responses, conduction speed, relative slowing and conduction failure were extracted. The baseline conduction speed was calculated from the initial latency, defined as the time delay between the stimulus and the induced response at the recording site, at the beginning of the stimulation. The conduction speed was then obtained by dividing the distance by the initial latency. The distance represents the hypothetical path taken by the axon from the recording site to the stimulation site and was approximated to be 2000 µm for all data points, based one the theoretical measurement of shortest possible path and longest possible path distances (Figure S1; Supporting Information). We assessed conduction failure by identifying the final stimulation pulse that elicited a detectable response. The ratio of the time until the final elicited response over the total duration of stimulation defined the “point of conduction failure”. We characterized the slowdown in conduction by calculating the relative slowing as follows:

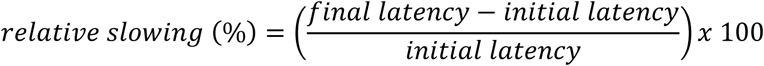

### Statistical analysis

All data points in the line plots represent the mean across all axon bundle mean individual induced axon responses per condition. All data are shown as mean ± SE of means, as noted in the figure legends. To summarize and visualize the distribution of conduction properties across conditions, boxplots were generated using the Seaborn library in Python. The boxplots display the median, interquartile range (IQR), and potential outliers for each group. The box represents the IQR (25th to 75th percentile), with the horizontal line inside indicating the median. Whiskers extend to 1.5 times the IQR, and data points beyond this range are plotted individually as potential outliers. All individual bundle data points are overlayed on the boxplots. Each data point in the boxplot represent the mean across induced axon responses per stimulated axon bundle. The Welch’s t-test was chosen to be the most appropriate choice for the data given the unequal variances and approximately normal distributions in both groups. Statistical analysis was done using Python. *** indicates p *<*0.001.

## Acknowledgements

The authors thank Silvio Scherr from the ETH Zurich Physics Workshop for laser cutting of the PMMA spacers to crosslink the gels. We are grateful to Dr Marios Tringides for reading the manuscript and providing valuable edits. The authors also gratefully acknowledge ScopeM for their support & assistance in this work, in particular Stephan Handschin for preparing and acquiring the TEM images. This work was supported by an ETH Zurich Postdoctoral Fellowship, for Tringides. ETH Zurich is acknowledged for financial support of the experiments, for both the MWT and JV. JV was additionally supported by Swiss National Science Foundation (SNF) and the Human Frontiers for Science Program. LS was supported by a grant by the SNF (310030_192075).

## Data availability

The data that support the findings of this study are available in a public repository at the time of publication.

