## supplemental data for "HydroMEA: A 3D Hydrogel Based Microfluidic Device to Study Electrophysiology for Myelinated Nerve-on-Chip"

### SUPPLEMENTAL INFORMATION

#### Materials and methods:

***Synthesis of alginate***: To prepare the nanoporous and porous alginate hydrogels, previously reported protocols were followed[68,69]. Briefly, alginate with two different molecular weights was used to form gels with ‘elastic’ or ‘viscoelastic’ properties, respectively: high Molecular Weight (HMW) alginate Protonal LF10/60, and Low Molecular Weight (LMW) alginate Protonal LF10/60 3mRad irradiation were functionalized with the ligand RGD, following previously published protocols[54,70,71].

***Preparation of alginate hydrogels:*** After sterilization and preparation of the RGD-alginate material, the powder was weighed and dissolved in sterile media at the desired weight/volume percentage (typically 2-2.5% w/v). The powder was left to fully dissolve in the media overnight, with a sterile stir bar in a sterilized scintillation vial. A sterile calcium sulfate solution (40 mM) was prepared, and diluted 5x in media. Formulations were either crosslinked with the desired amount of calcium, using a double-barrel syringe method connected by a Luer lock, as previously described[26,72,73], or by casting the precursor solution, freezing the material in a tissue culture plate (typically 12, 24, or 48 well plates) before being frozen at -20° C. Once frozen, the plates were transferred to a lyophilizer (Freezone, Labconco). Gels were cross-linked by ionic crosslinking with Ca2+ dissolved in ethanol at a concentration of 450mM, as previously reported.

***PEG-based hydrogel fabrication***[74]: A polymer precursor solution was prepared by combining 13 µL of 4-arm 20 kDa PEG-norbornene (PEG-NB, 25% w/v) with 1 µL of MMP-cleavable peptide linker (KCGPQG↓IWGQCK, Genscript) (final concentration: 2.35 mM) and 1.74 µL of cell adhesion peptides (CRGDS or CDPGYIGSR, Genscript) (final concentration: 1.8 mM). To provide additional cell-binding motifs necessary for neural cell attachment and growth, 5.5 µL of laminin was incorporated into the formulation. Since the MMP-cleavable linker creates an acidic environment, 0.3 µL of NaOH was added to normalize the pH to approximately 7.4, ensuring optimal conditions for cell viability. The prepared neural cell suspension (76.96 µL) was then carefully mixed with the polymer precursor solution to achieve a final cell density of 2 × 106 cells/mL. Immediately prior to photocrosslinking, 2.5 µL of the photoinitiator lithium phenyl-2,4,6-trimethyl-benzoylphosphinate (LAP, 5 w/v) was added to the mixture, bringing the total hydrogel precursor volume to 100 µL. The complete hydrogel precursor solution was gently pipetted between two glass slides pre-treated with Sigmacote® (Sigma, SL2) to ensure non-adhesion to the surfaces. The solution was then crosslinked via photoinitiated thiol-ene chemistry, forming covalent bonds between the thiol groups of the MMP-cleavable linkers and the norbornene groups of PEG-NB. Crosslinking was achieved through exposure to blue light (λ = 405 nm, I = 14.5 mW cm−2) for 90 seconds, resulting in a mechanically stable hydrogel with encapsulated neural cells.

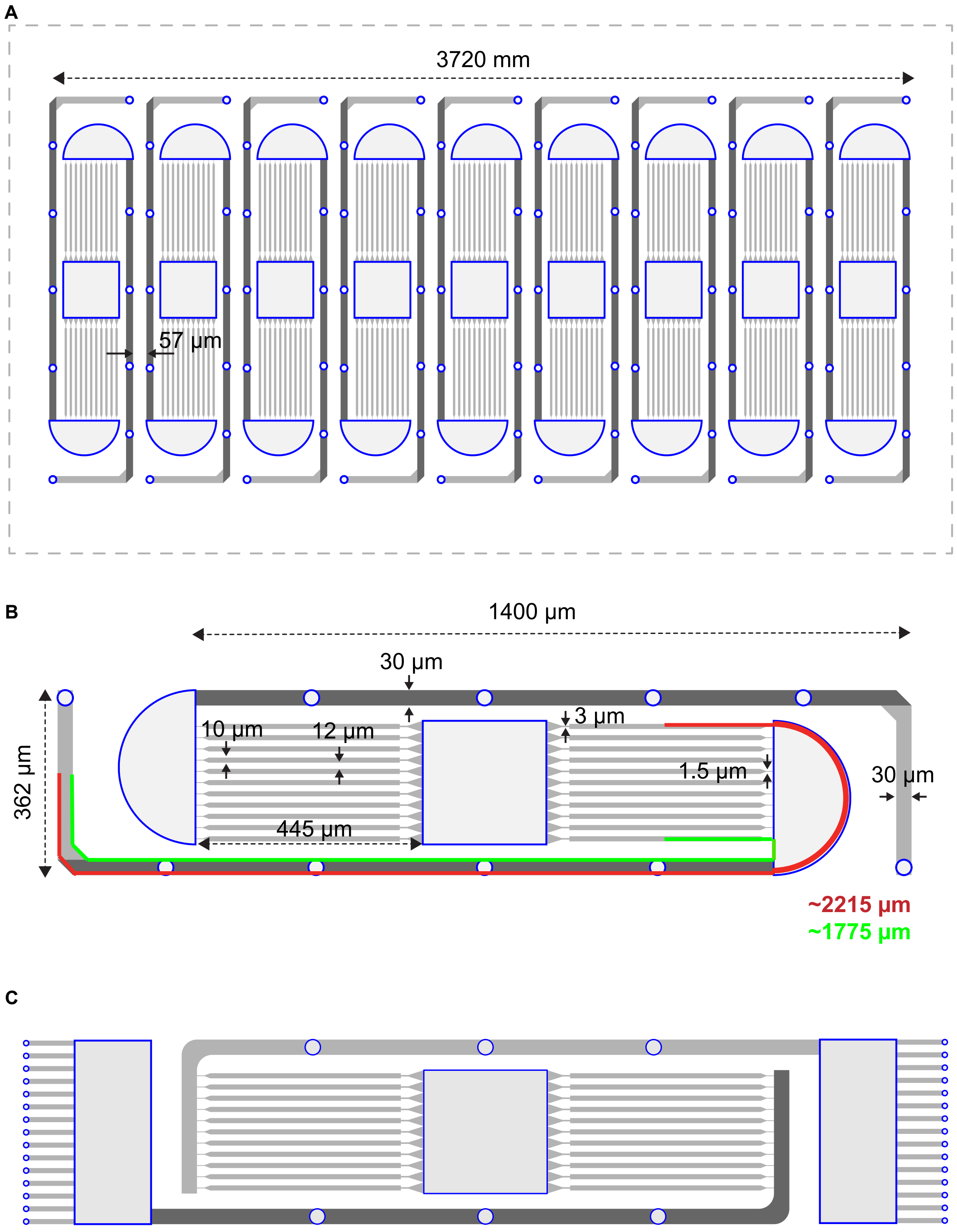


**Figure S1: Detailed network design dimensions**. **A**: Nine parallel networks fit on the sensing area of the HD-MEA. **B**: Dimension of the features of a single network. The theoretical shortest and longest paths from the stimulation site and recording side are indicated with pink and red lines. The average of both distances is 2 mm and used in the calculation of the conduction speed. **C**: A slightly different design of B was used in **Figure 5**. The main design difference is the location of the Schwann cell seeding wells. In this case they are located at the end of the axon bundle channels instead of at the beginning, between microchannels and wide channel. While this design does not allow stimulation at the end of the axon bundle channel, it could be used as well for optimizing hydrogel integration on glass and be used for optical microscopy-based control.

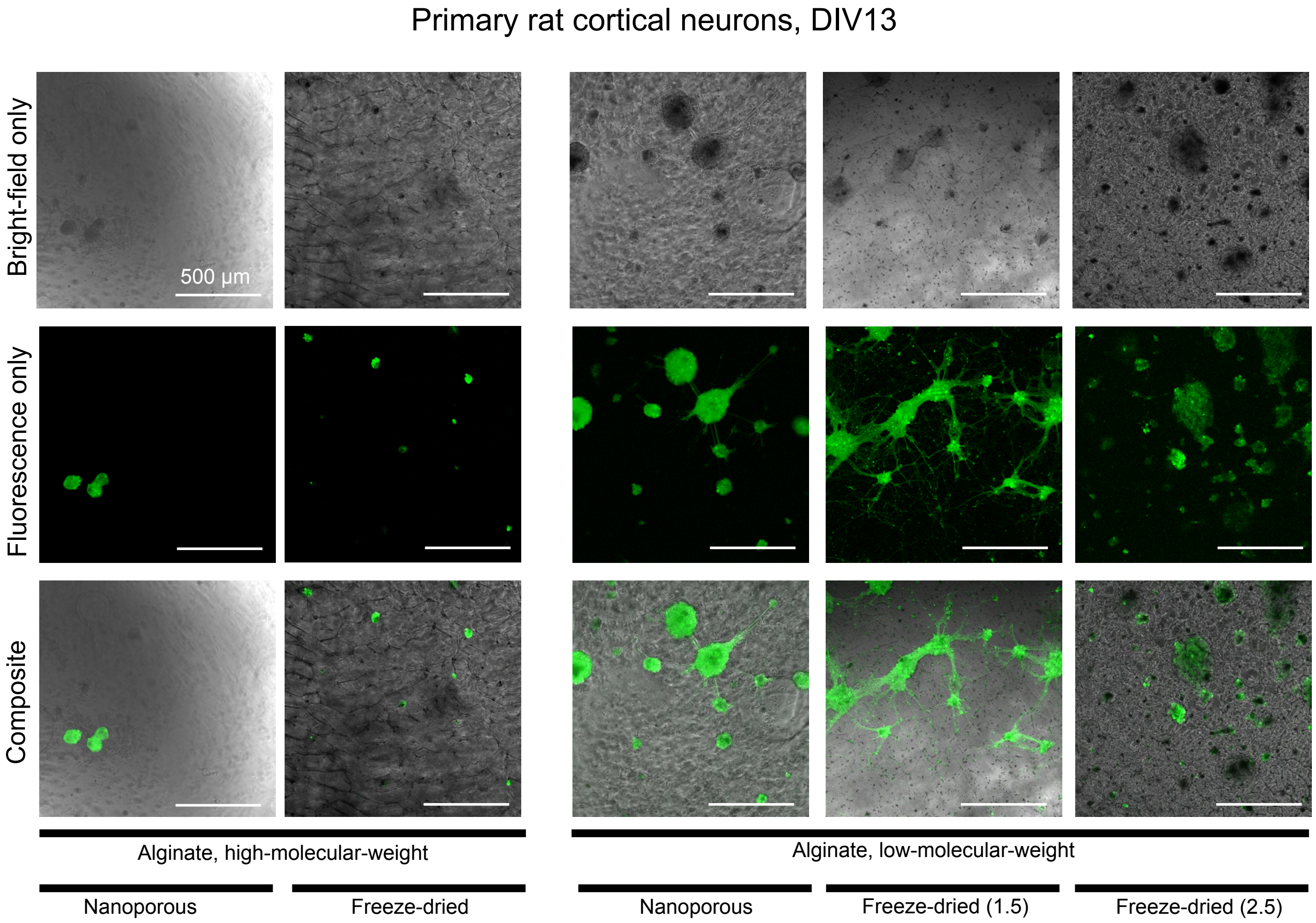


**Figure S2:** Feasibility of growing neurons in alginate hydrogel. Photomicrographs (bright-field only, fluorescence- only and composite) showing viability and growth of primary rat cortical neurons in different alginate hydrogel formulations after 13 days in vitro (DIV). Formulations included the use of low-molecular-weight and high-molecular-weight alginate material, to form nanoporous and porous (freeze-dried) 3D scaffolds. Live neurons were stained with Calcein-AM. Scale bar = 500 µm.

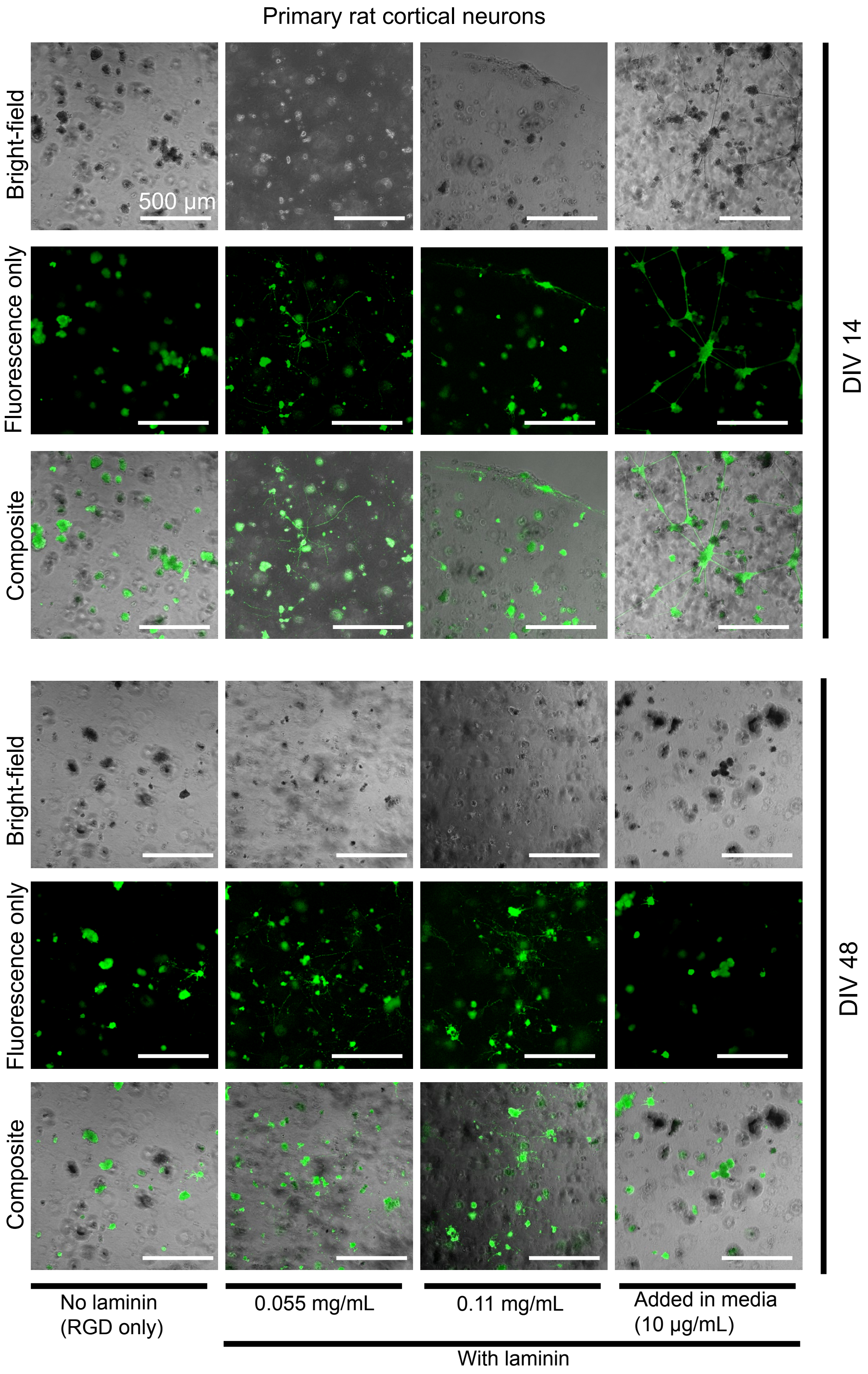


**Figure S3: Feasibility of growing neurons in PEG-based hydrogel.** Photomicrographs (bright-field only, fluorescence-only and composite) showing viability and growth of primary rat cortical neurons in PEG-based hydrogel with different concentrations of laminin, at DIV 14 and DIV 48. Neurite outgrowth was enabled in conditions with laminin, and promoted with higher concentrations when supplemented in hydrogel formulation directly. Live neurons were stained with Calcein-AM. Scale bar = 500 µm.

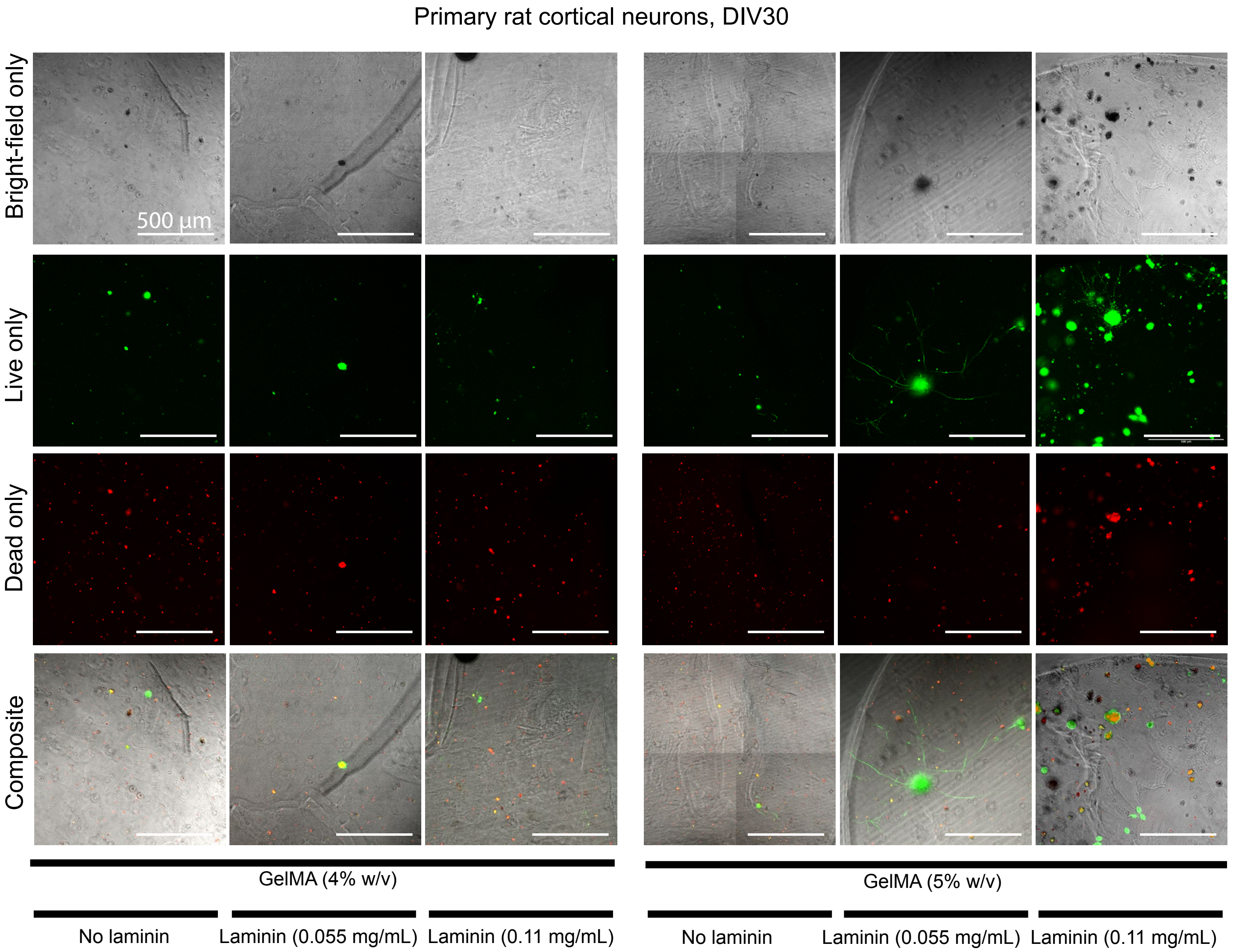


**Figure S4: Feasibility of growing neurons in GelMA-based hydrogel.** Photomicrographs (bright-field only, fluorescence-only for live/dead staining, and a composite image overlay) showing viability and growth of primary rat cortical neurons in GelMA hydrogel with different concentrations of GelMA and laminin, at DIV 30. Neuron viability and neurite outgrowth were promoted for 5% GelMA in presence of laminin. Live neurons were stained with Calcein-AM and dead neurons were stained with ethidium-homodimer-1. Scale bar = 500 µm.

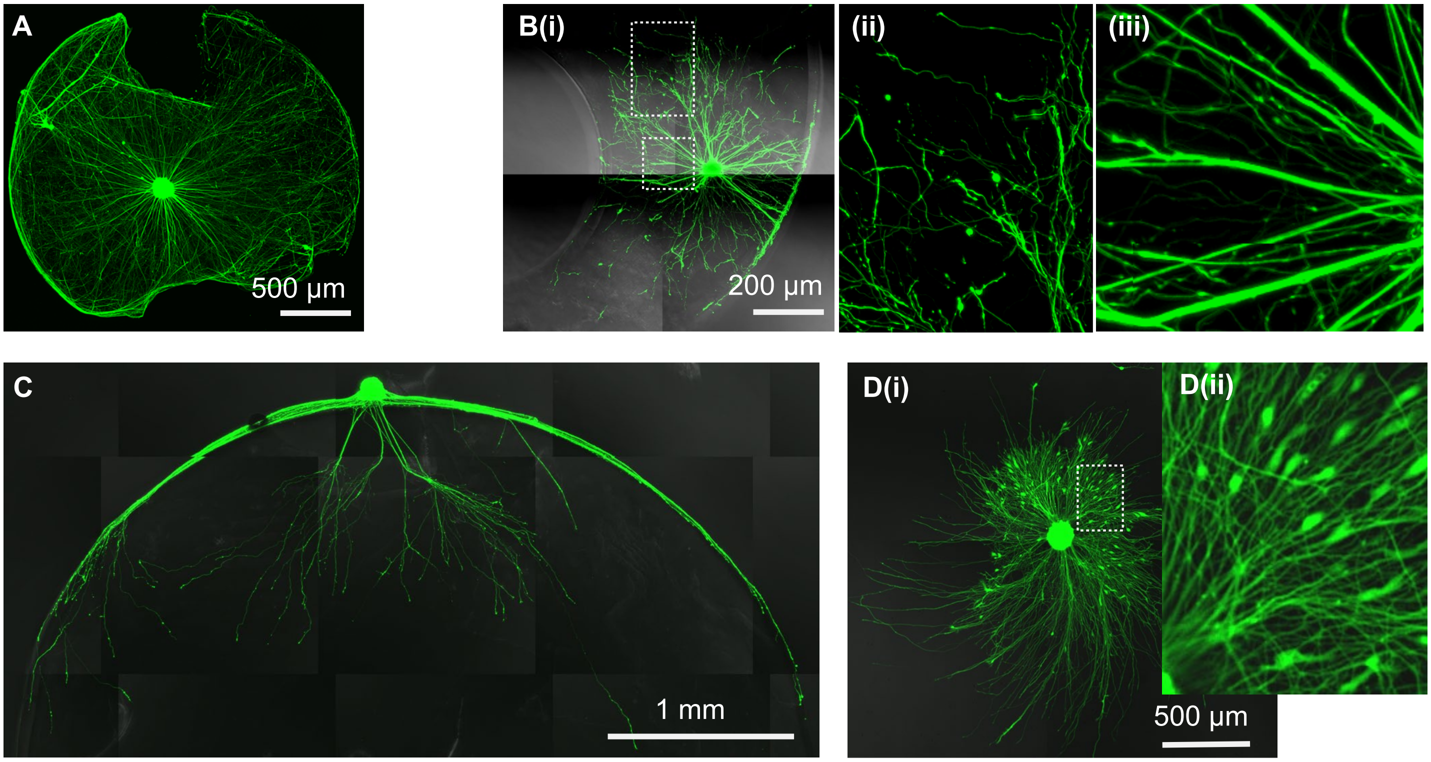


**Figure S5: Comparison of neurite outgrowth behavior depending on the substrate modalities (2D versus 3D)**. **A**. Z maximum intensity projection of the different z planes shown in main Figure 3 for a bulk gel in which one neuron spheroid grew axons. **B.** Another bulk gel supported 3D growth. (i) zoom on the nonlinear growth of axons inside the gel. (ii) Zoom on the bundle-like growth of axons before penetrating the gel. **C.** Example of a spheroid growing from the surface of the gel towards the inside. **D.** Neurite outgrowth from a spheroid on a 2D coated-glass substrate. Axons grow in a straighter manner than in 3D and cell bodies migrate outside of the spheroid core unlike in 3D.

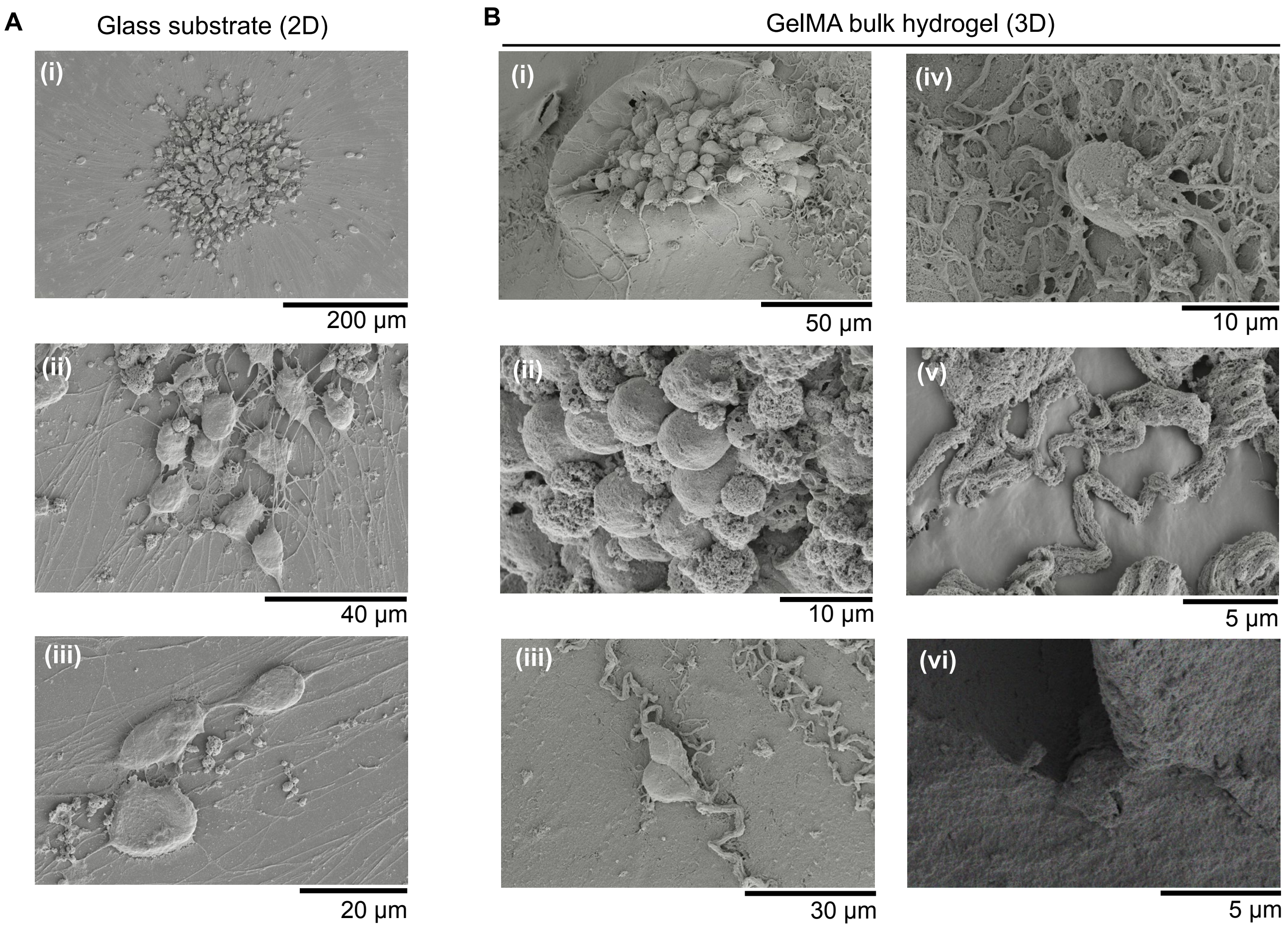


**Figure S6: Scanning Electron Microscopy (SEM) of 3D versus 2D grown hSNs**. **A:** SEM photomicrographs of neurons grown on 2D substrate (laminin-coated glass). (i): Neurons were seeded as a spheroid, and cell bodies migrated out, comparably to the fluorescence photomicrographs shown in Figure S5. (ii-iii): Cell bodies have a flattened morphology and neurites grow in a straight path manner on the substrate. **B:** SEM photomicrographs of neurons grown in GelMA-based hydrogel (3D). (i): Neurons were seeded as a spheroid, with cell bodies staying in the spheroid core. (ii) Cell bodies have a more round morphology compared when grown in 2D. (iii-v): Different zoom levels showing axons grown in a meandering manner. (vi): GelMA hydrogel exhibits porosity on the micron-scale.

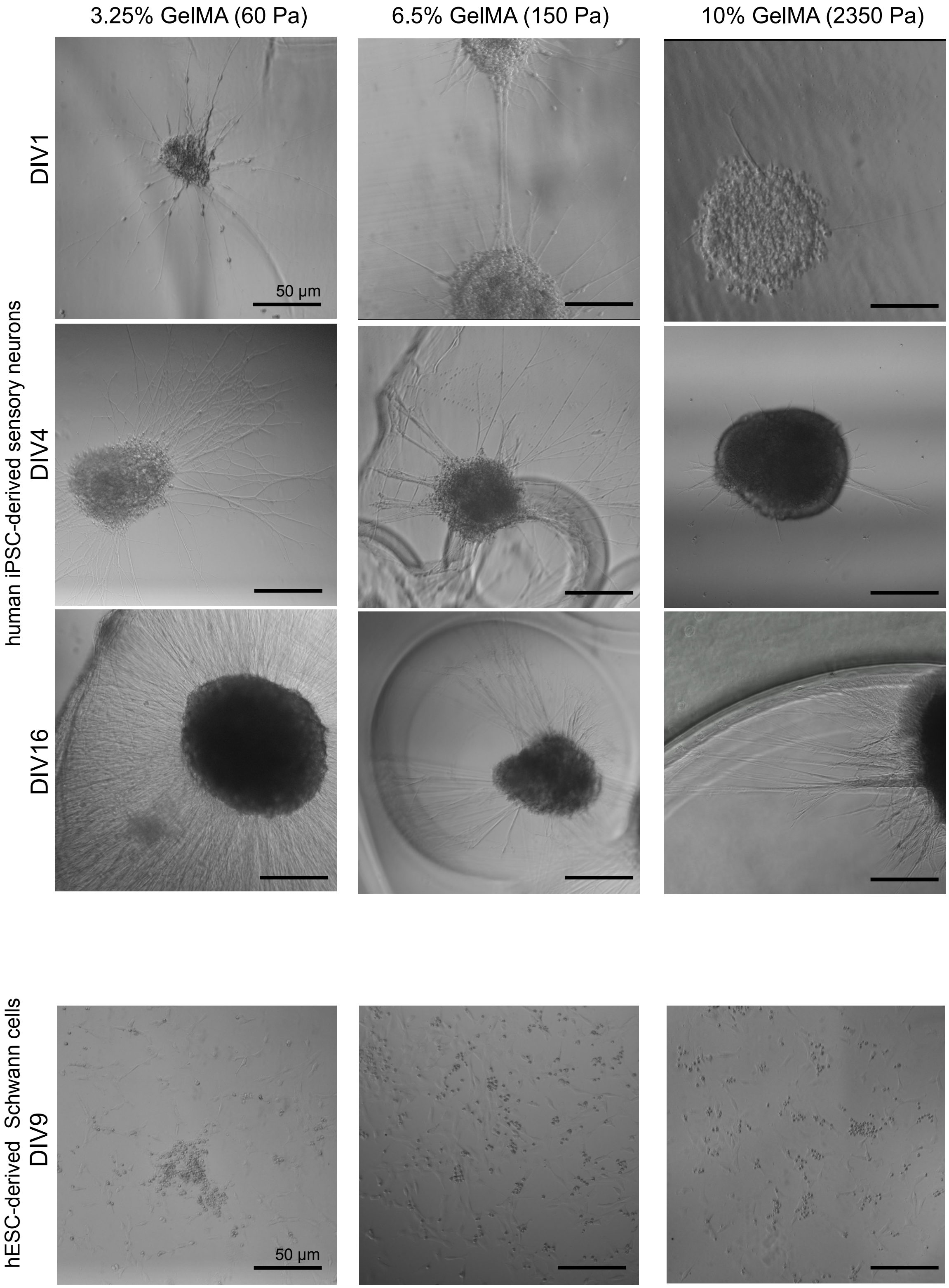


**Figure S7: Stiffness effect on hiPSC-derived sensory neurons (hSN) and hESC-derived Schwann cells.** Representative bright-field photomicrographs showing neurite outgrowth patterns of encapsulated spheroids at different DIV for three different stiffness. Neurites tend to grow more and faster inside the softer bulk gels. Schwann cells seem to be viable and proliferate in a similar manner regardless of the stiffness of the gel. Scale bar = 50 µm.

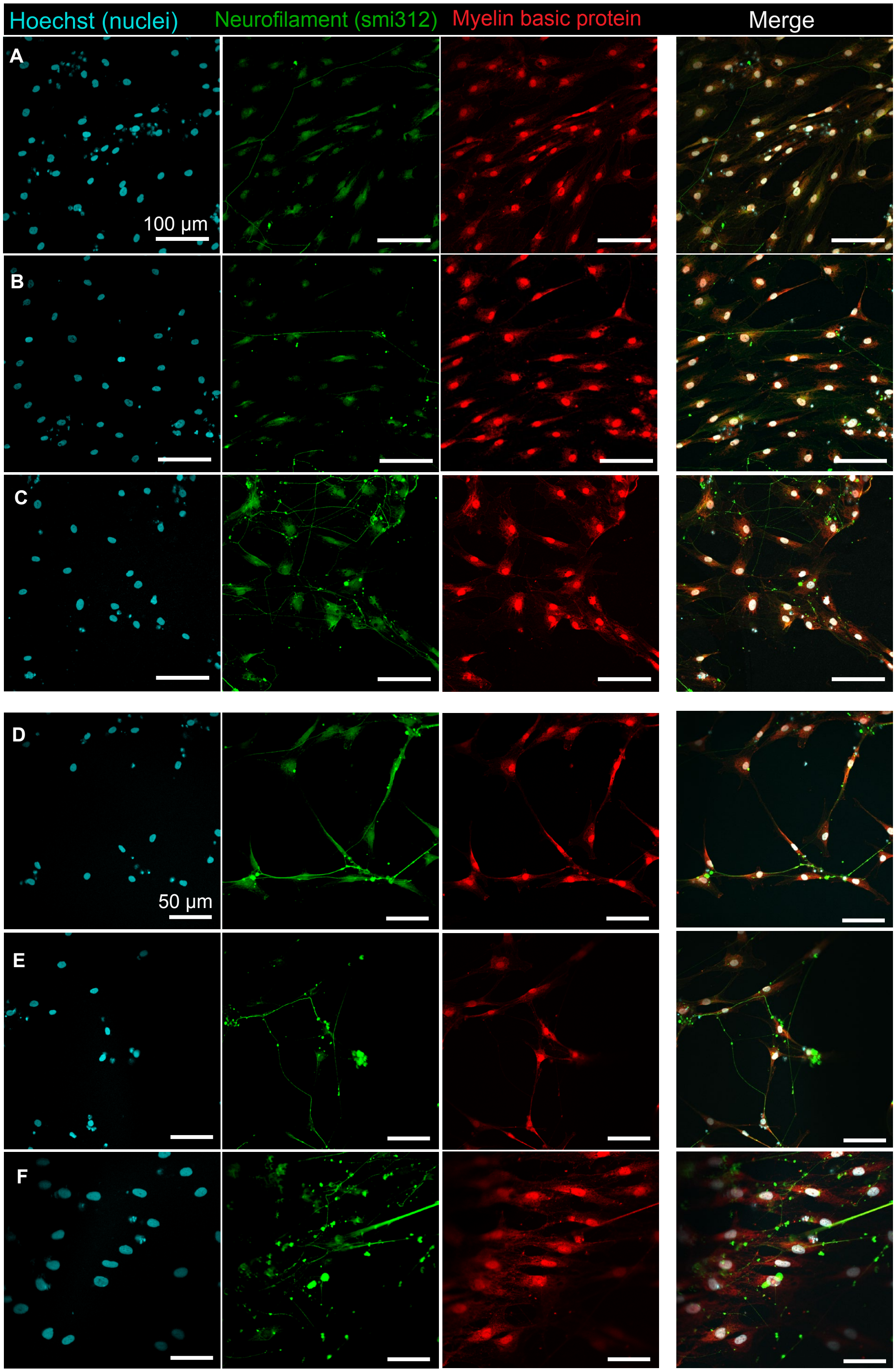


**Figure S8: Representative immunofluorescence photomicrographs of GelMA bulk hydrogel supporting long-term viability of co-cultures of hSNs and hSCs (DIV56).** Schwann cells interact with axons in a selective way. Photomicrographs in **A-C** show some Schwann cell-free axons among Schwann cells while **D-F** show Schwann cell-axon contacting and alignment scenarios. Scale bar = 100 µm (A-C) and 50 µm (D-F)

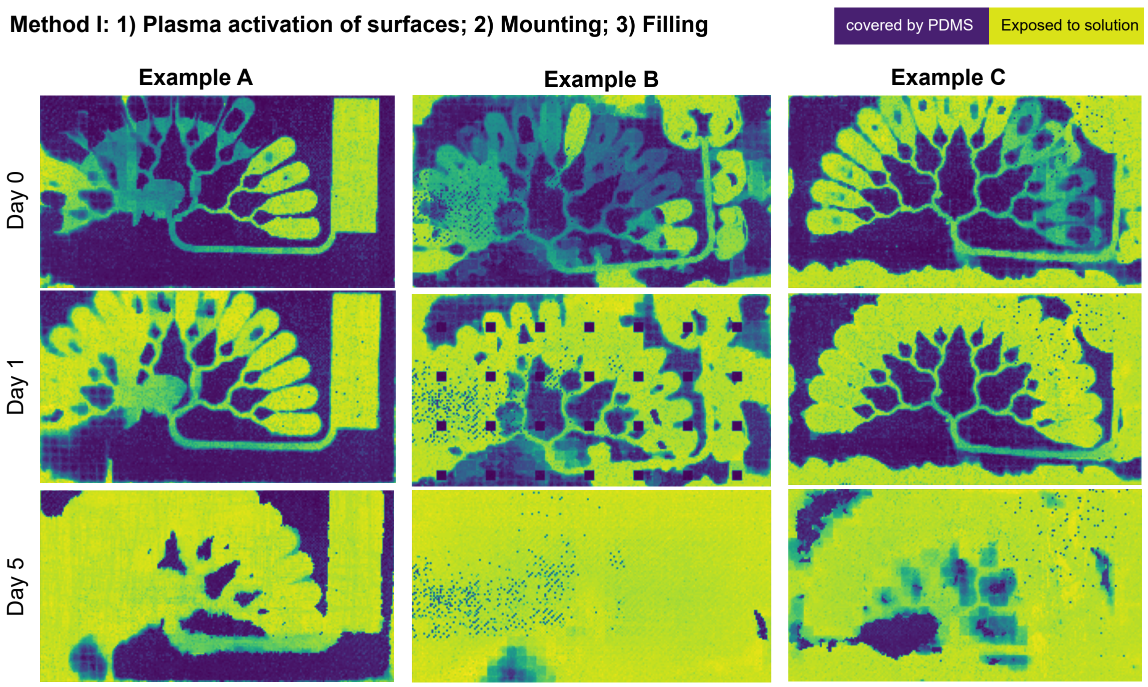


**Figure S9: Impedance maps detect PDMS microstructure on the HD-MEA with Method I, insufficient for sustained attachment.** Impedance maps reveal the liquid solution underflow below PDMS microstructure placed on top of a HD-MEA over days when placing PDMS after performing plasma cleaning of both surfaces (bottom face of PDMS microstructure and MEA substrate.

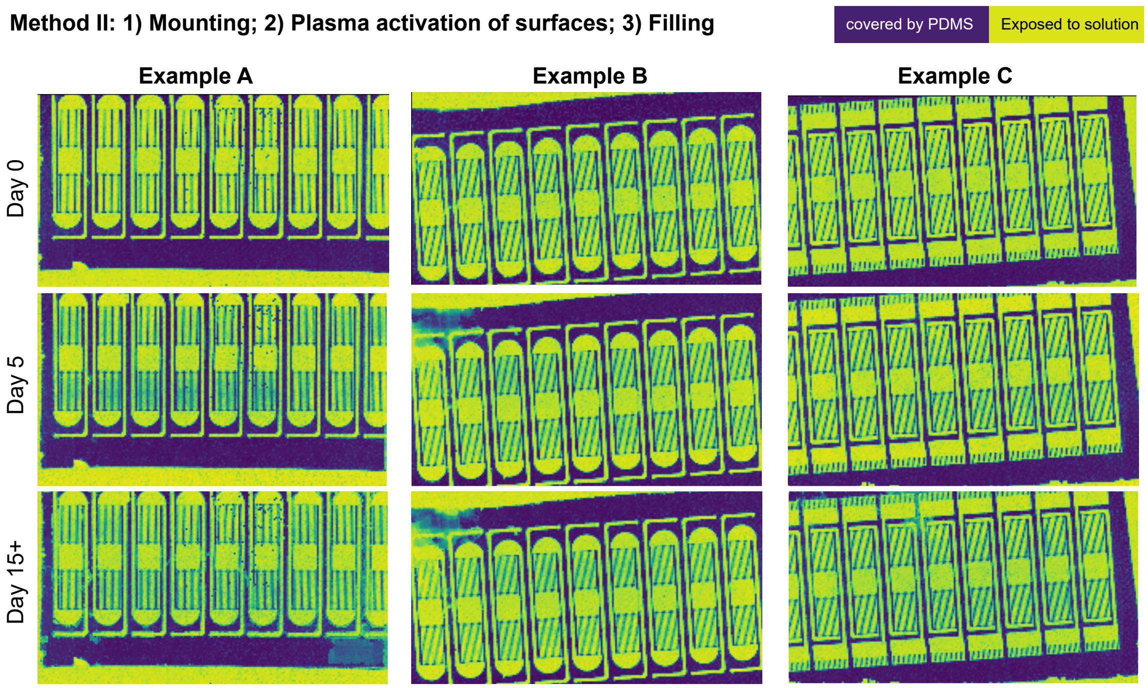


**Figure S10: Impedance maps detect PDMS microstructure on the HD-MEA with Method II, sufficient for sustained attachment.** Impedance maps revealing stability of PDMS microstructure adhesion placed on CMOS MEA over days when placing PDMS on CMOS MEA before performing plasma cleaning of the pre-mounted construct.

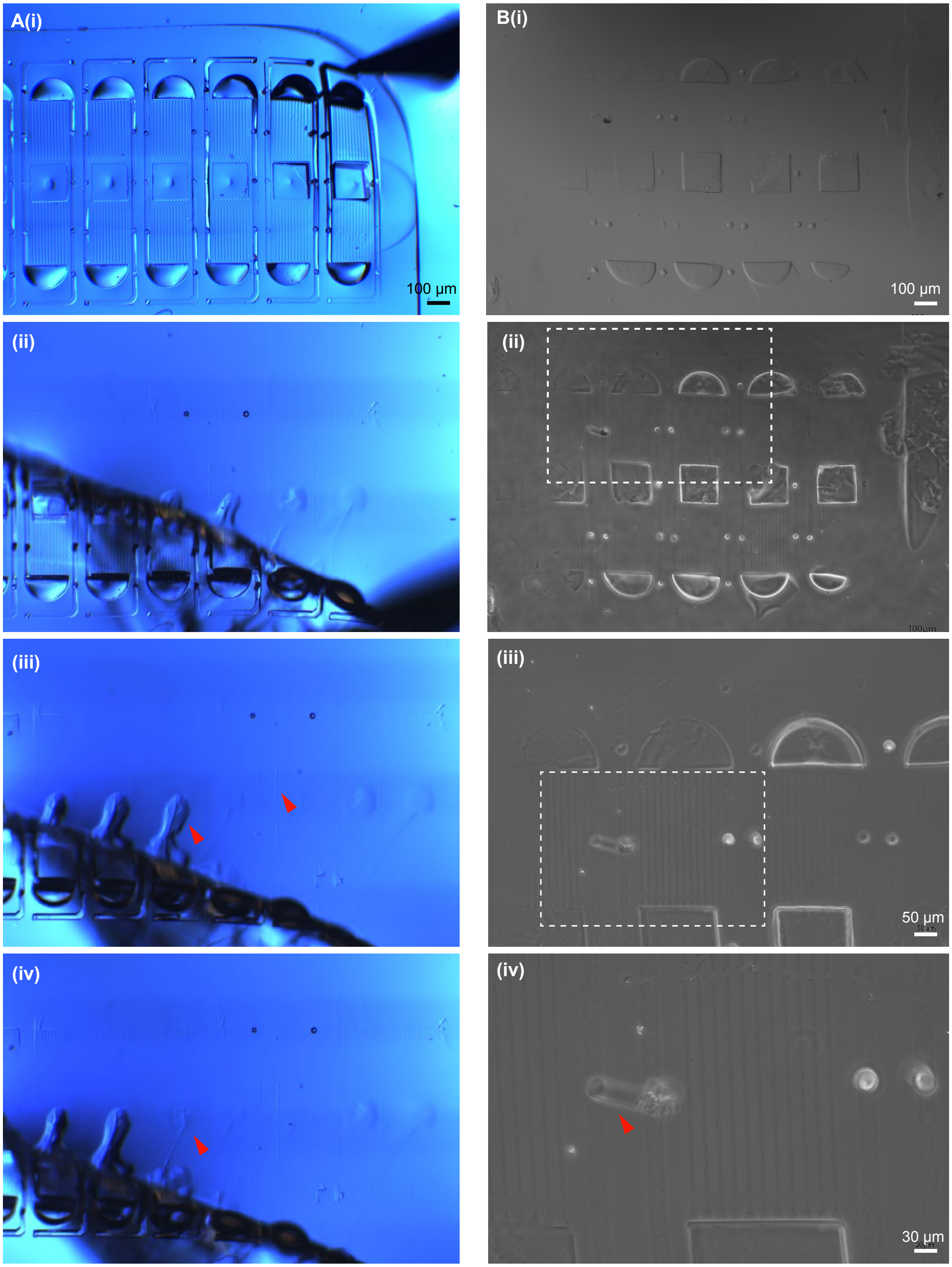


**Figure S11: GelMA hydrogel residues inside PDMS microstructure microchannels upon peel-off. A:** A PDMS microstructure was lifted with tweezers (right corner in (i) after cross-linking the GelMA hydrogel precursor solution to ensure the gelling inside microchannels (i.e.; right red arrow in (iii). (i-iv) show different moments during the peel-off. It is worth mentioning that residues of GelMA gel might get stuck inside the PDMS microstructure and not only on the glass substrate during such procedure as illustrated by the transition between (iii) and (iv) and indicated by red arrow. **B:** Additional example of a different microstructure showing left over hydrogel traces revealing microchannel design. (iv) indicates where the gel negative of one of the diffusion wells.

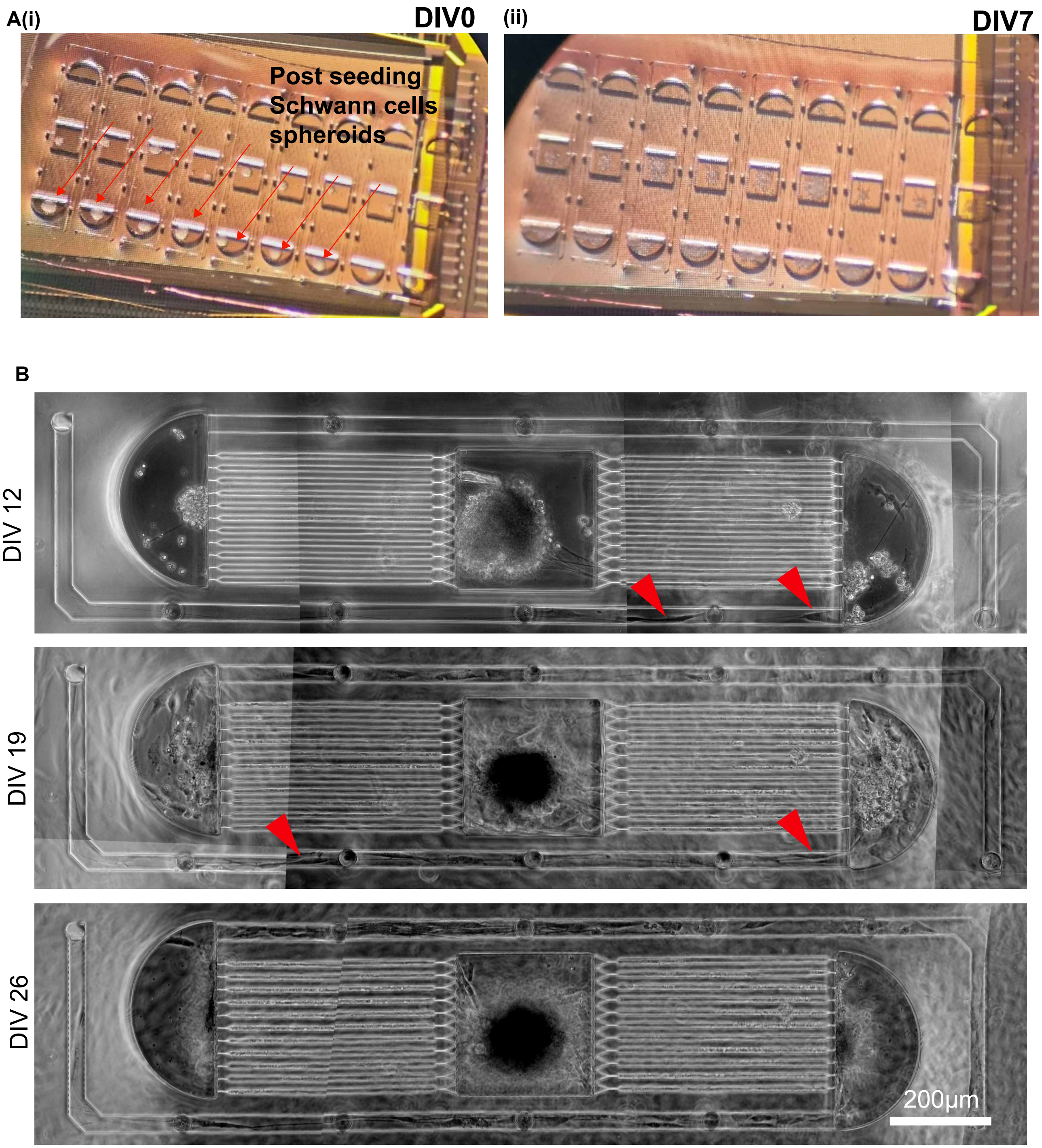


**Figure S12: Overview of the platform and Schwann cell placement at early stages A:** Photo taking through a stereomicroscope of the PDMS microstructure on HD-MEA (i) after seeding hSC spheroids and (ii) one week later. **B:** Bright-field photomicrograph of one example network showing the progressive migration of hSC from the lateral compartment towards the bundle channel for DIV 12, DIV 19 and DIV 26. Red arrows highlight hSC migration into the nerve-forming compartment.

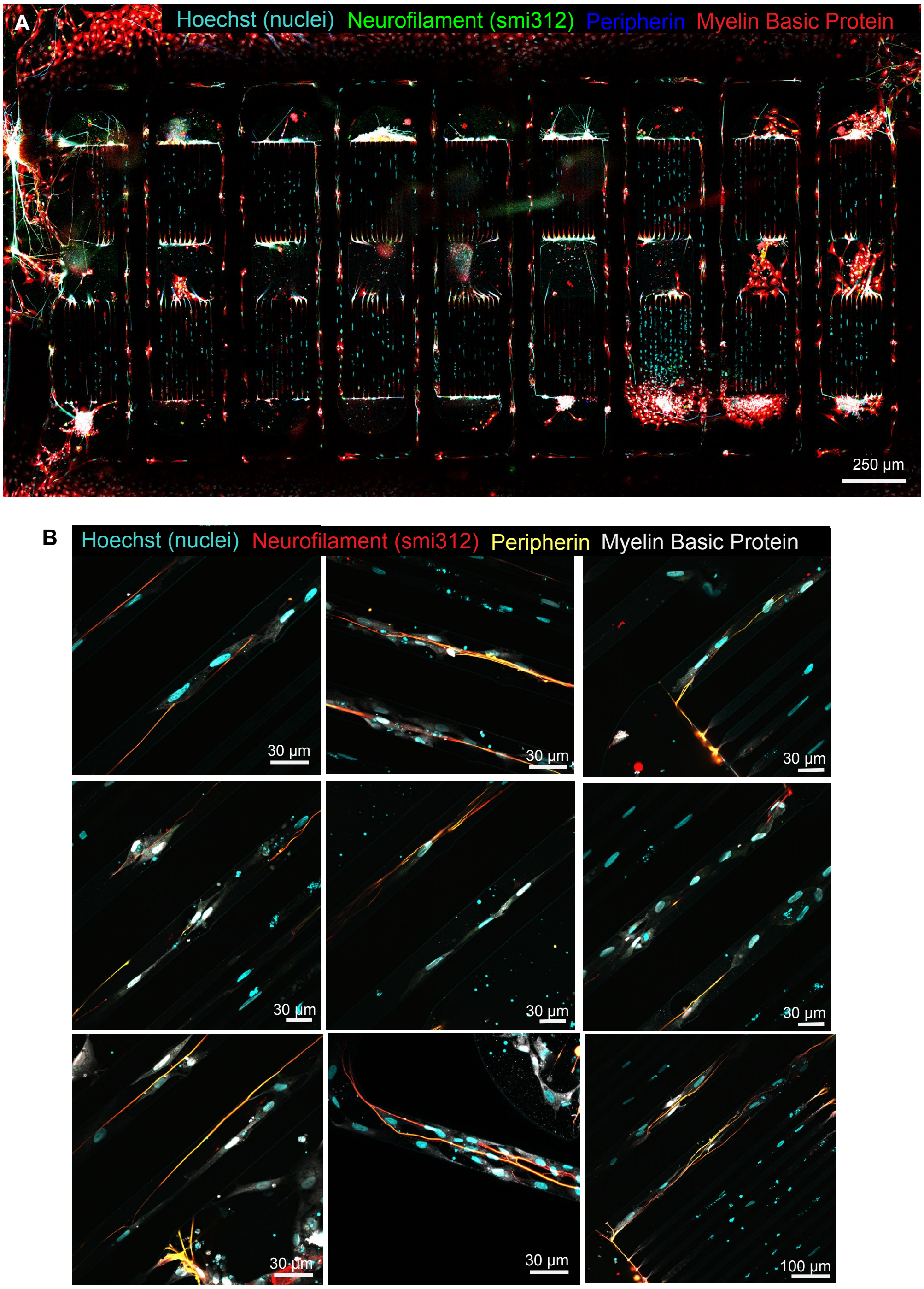


**Figure S13: Representative immunofluorescence photomicrographs of the integrated co- culture in microstructure. A**: Example of 9 distinct networks on glass, exhibiting variability in cell arrangement and migration across networks. Sample was fixed at DIV 56. Scale bar = 250 µm. **B:** Additional zoomed-in images of the axonal segments of hSNs with hSCs in microchannels showing interactions between the two cell types.

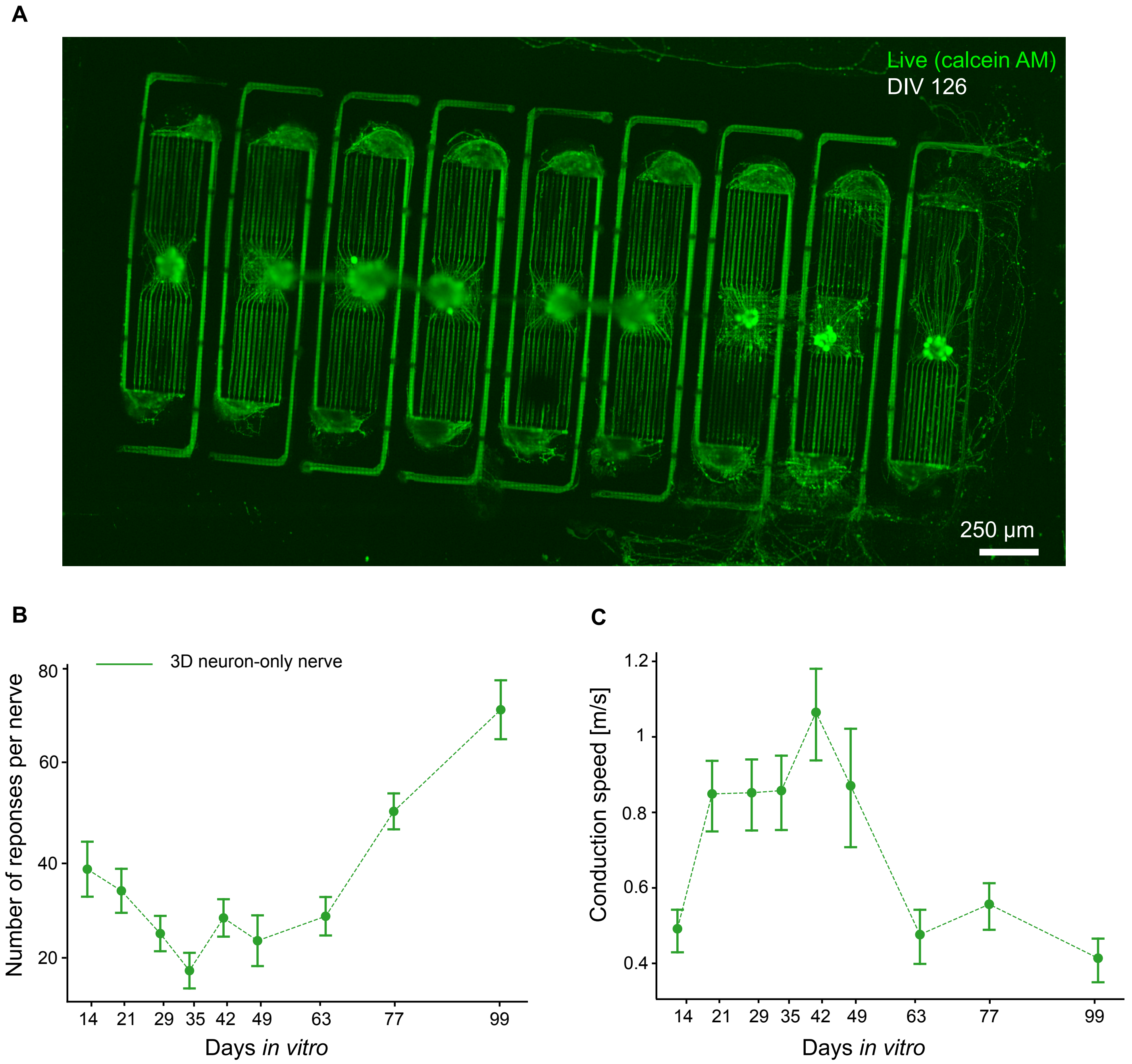


**Figure S14: Feasibility of long-term 3D nerve growth and electrophysiology on HD-MEA with hSN only.** **A:** Photomicrograph of fluorescently labeled live (calcein AM, green) neurons at DIV 126, at the final timepoint of electrophysiology measurements. **B:** Number of recorded responses per stimulated nerve over 100 DIV, averaged over 9 independent networks. **C:** Conduction speed over 100 days in vitro. N = 16 nerves per time point. Mean and standard error are plotted for each graph.

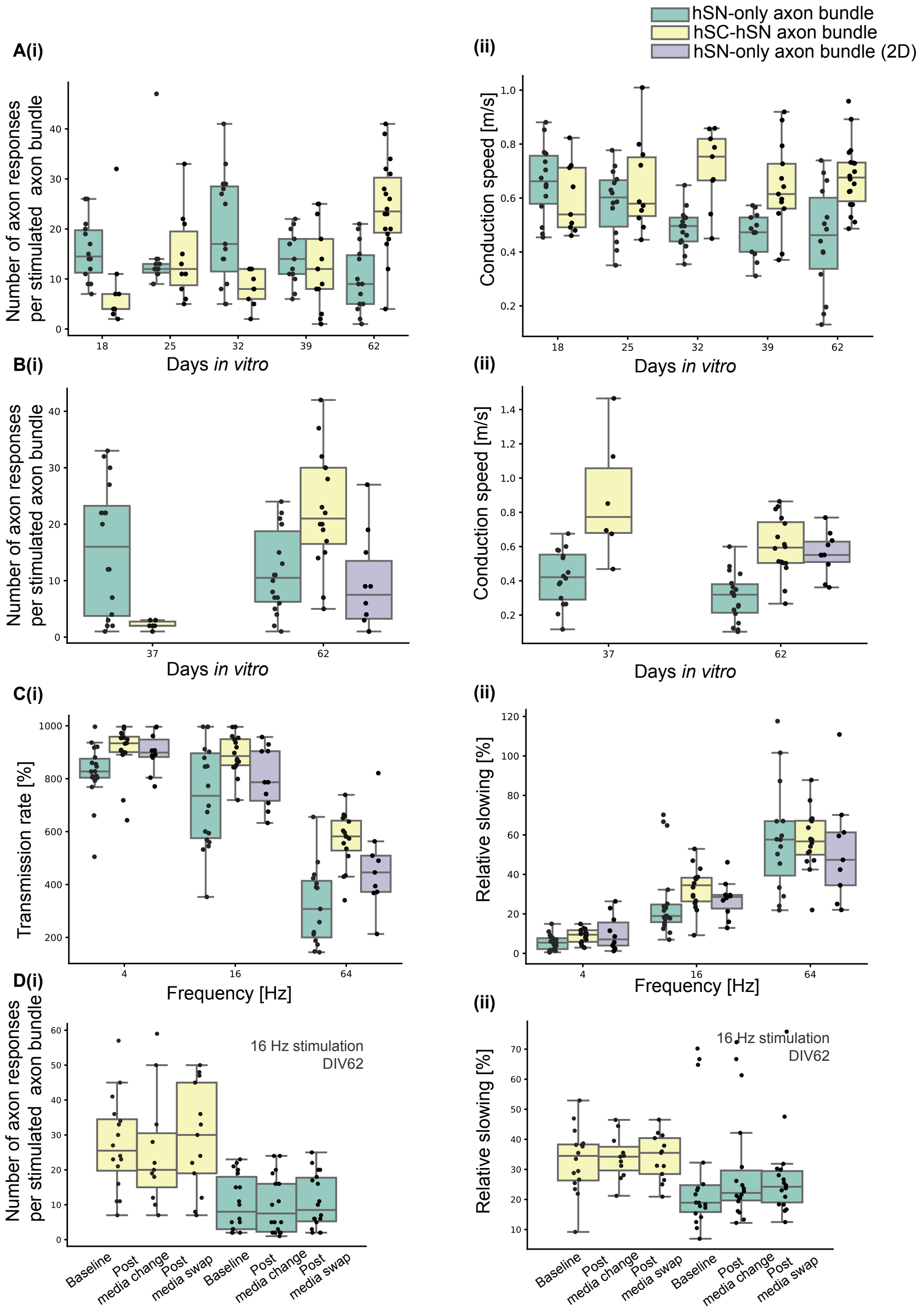


**Figure S15: Electrophysiological characterization of hydroMEA. A**(i) Boxplot representation of the line plot for the (i) number of axon responses and (ii) conduction speed of stimulated axon bundles across DIV shown in main Figure 7C(i) and (ii). **B:** Supplementary data from a different experiment showing (i) number of axon responses and (ii) conduction speed for DIV 37 and DIV 62 with an added "2D" condition at DIV 62. **C:** Boxplot representation of the line plots of (i) transmission rate and (ii) relative slowing across stimulation frequencies shown in main Figure D(i) and (ii). **D:** Boxplots for (i) number of axon responses per stimulated axon bundle and (ii) relative slowing for the different media change conditions.

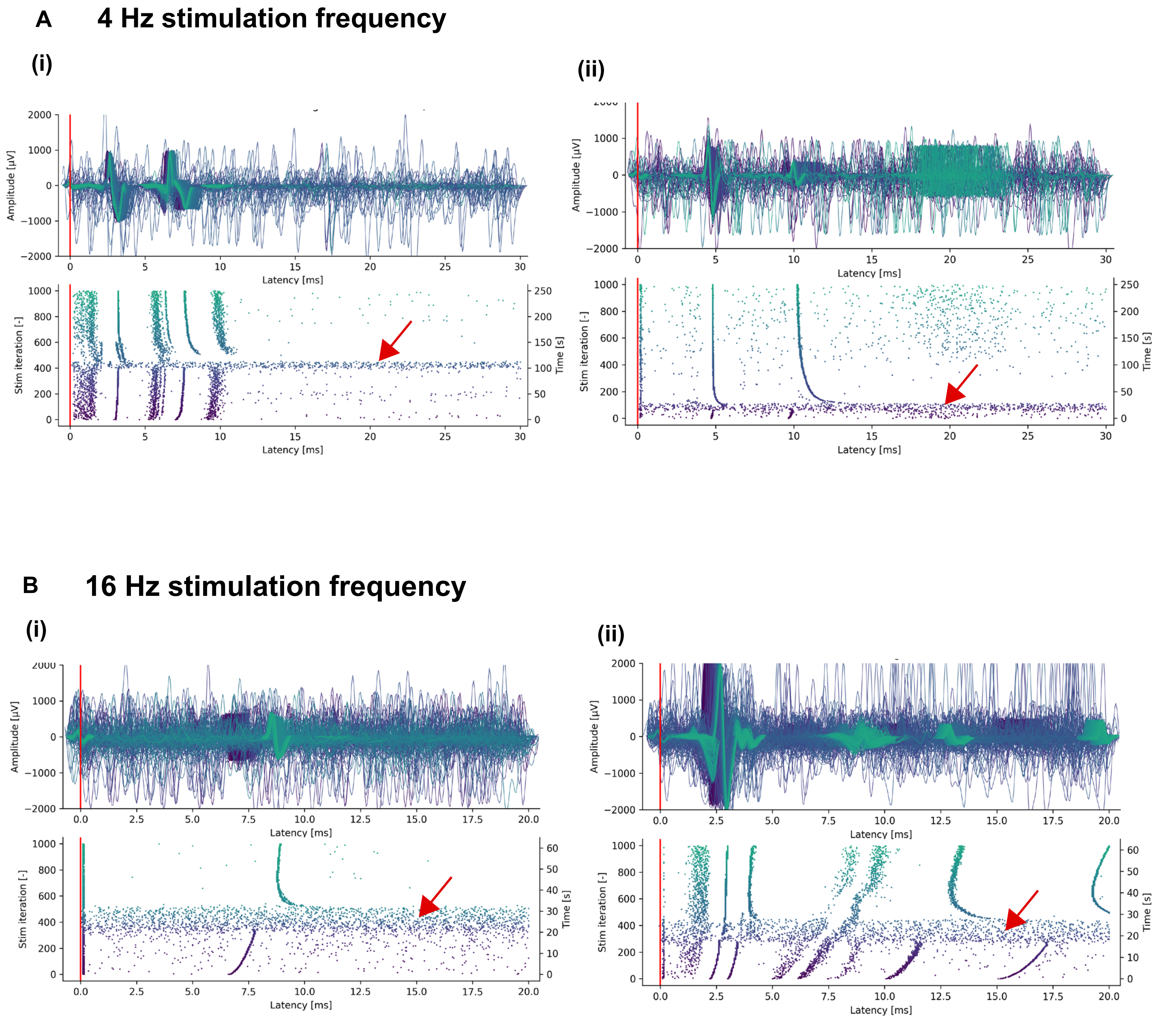


**Figure S16: Post-stimulus raster plot examples showing cases of transient increase in spontaneous activity specifically observed in the hSC-hSN axon bundles. A**: Two independent axon bundle examples stimulated at 4 Hz. The sudden increase in spontaneous activity is visible from the horizontal "band" showing denser amount of detected spike events, lasting about ten minutes and impeding stimulation-induced activity, and inducing an interruption in the response (vertical "bands"). The same effect happens in (ii) but earlier in the stimulation train, suggesting that the timing of this transient activity is not related to the onset of the stimulation. **B:** Similar to A for 16 Hz stimulation, which highlight the change in axonal response before and after this occurs. Indeed, responses that exhibit a stronger activity-dependent slowing seem to re-start from later latencies, and tend to go back towards baseline latency, or initial relative slowing rate. The duration of the event confirms the one in **(A)** and lasts for 10 minutes.
